# Metabarcoding faecal samples to investigate spatiotemporal variation in the diet of the endangered Westland petrel (*Procellaria westlandica*)

**DOI:** 10.1101/2020.10.30.360289

**Authors:** Marina Querejeta, Marie-Caroline Lefort, Vincent Bretagnolle, Stéphane Boyer

## Abstract

As top predators, seabirds can be indirectly impacted by climate variability and commercial fishing activities through changes in marine communities. However, high mobility and foraging behaviour enables seabirds to exploit prey distributed patchily in time and space. This capacity to adapt to environmental change can be described through the study of their diet. Traditionally, the diet of seabirds is assessed through the morphological identification of prey remains in regurgitates. This sampling method is invasive for the bird and limited in terms of taxonomic resolution. However, the recent progress in DNA-based approaches is now providing a non-invasive means to more comprehensively and accurately characterize animal diets. Here, we used a non-invasive metabarcoding approach to characterize the diet of the Westland petrel (*Procellaria westlandica*), which is an endangered burrowing species, endemic to the South Island of New Zealand. We collected 99 fresh faecal samples at two different seasons and in two different sub-colonies. Our aims were to describe the diet of the Westland petrel, investigate seasonal and spatial variation in the petrels’ diet, and assess potential impacts of the New Zealand fishery industry. We found that amphipods were the most common prey, followed by cephalopods and fish. Our results could be the result of natural foraging behaviour, but also suggest a close link between the composition of prey items and New Zealand’s commercial fishing activities. In particular, the high abundance of amphipods could be the result of Westland petrels feeding on discarded fisheries waste (fish guts). Our results also showed significant differences in diet between seasons (before hatching vs chick-rearing season) and between sampling sites (two sub-colonies 1.5 km apart), indicating plasticity in the foraging strategy of the Westland petrel. Due to its non-invasive nature, metabarcoding of faecal samples can be applied to large numbers of samples to help describe dietary variation in seabirds and indicate their ecological requirements. In our example, dietary DNA (dDNA) provided valuable information regarding the dietary preferences of an iconic species in New Zealand’s unique biodiversity. dDNA can thus inform the conservation of endangered or at-risk species that have elusive foraging behaviours.

## Introduction

The study of animal diets is a critical component in many aspects of ecology, including community ecology (Corse et al., 2010), population dynamics (Morrison et al., 2014; Read and Bowen, 2001) and conservation biology (Lyngdoh et al., 2014; Xiang et al., 2012). In predators, spatial and seasonal changes in diet composition may reflect a certain degree of flexibility in foraging behaviour (Whelan et al., 2000). This variation could be relevant for understanding trophic interactions and conserving endangered species (Davies et al., 2001; Farias and Kittlein, 2008; Vander Zanden et al., 2000; Vinson and Angradi, 2011). Shedding light on dietary patterns is essential in the case of seabirds, which are top predators within marine ecosystems.

Seabirds are known to modify their feeding habits depending on the time of the year (Harding et al., 2007; Kowalczyk et al., 2015) and their breeding site (McInnes et al., 2017a; Thompson et al., 1999). These birds spend most of their lives at sea but during the breeding season, some remain in coastal areas, as their foraging trips are restricted in number and length to allow them to regularly feed their chicks in the nest. To achieve this, seabirds have adopted a variety of foraging strategies (McInnes et al., 2017a; Ydenberg et al., 1994), such as switching between short and long foraging trips to feed their chicks while maintaining their body condition during the breeding season (Baduini, 2003; Ropert-Coudert et al., 2004), or providing the chicks with highly nutritive processed stomach oil (Baduini, 2003). The majority of studies that aim at describing the diet of seabirds have been carried out during the chick-rearing period only. Often, this is because data are collected based on the morphological analysis of regurgitates obtained from parents coming back to the nest to feed their chicks (Calixto-Albarrán and Osorno, 2000; Croxall et al., 1988; Klages and Cooper, 1992; Suryan et al., 2002). However, this approach considers prey communities as a fixed parameter across time, instead of treating it as a dynamic variable (Barrett et al., 2007; Komura et al., 2018).

Consequently, many studies do not explore diet plasticity, despite the ability to switch to new prey representing a potential mechanism to avoid large population declines that can lead to local extinctions of threatened populations (Marone et al., 2017). Many seabird populations have been decreasing rapidly in recent years (Grémillet et al., 2018; Thibault et al., 2019). Detailed spatiotemporal knowledge of their diet preferences is key to understanding and managing current and future threats, including commercial fishing activities or climate-driven changes to their ecosystem (Frainer et al., 2017).

Selecting the correct experimental design for diet analyses is challenging, especially in the case of seabirds, as it is mainly based on rare direct observations (Ocké, 2013). For decades, the morphological identification of stomach contents or regurgitates has been widely used to identify prey items of predators (Carreon-Martinez and Heath, 2010; Egeter et al., 2015; Freeman, 1998; Imber, 1976; Krüger et al., 2014). However, this methodology usually requires that gut content is obtained by stimulating regurgitation after capturing individual birds through a technique that has been called “lavage” (Barrett et al., 2007; Ryan and Jackson, 1986; Wilson, 1984). Such an invasive sampling method (Lefort et al., 2019) is not only unethical, but also potentially dangerous for the birds. Furthermore, the efficiency of this method is usually limited because many individuals would have empty stomachs while sampled, and highly digested prey items may not be identifiable to genus or species level.

The ability to identify prey remains from stomach content also varies in relation to prey species, because some species (in particular soft-bodied prey) are digested faster than others, making the taxonomic classification and species identification difficult or even impossible (Boyer et al., 2015; Deagle et al., 2007; Gales, 1988). As a consequence, soft-bodied animals are often overseen in prey biomass calculations, which could also lead to biases in the characterization and quantification of the diet. Other standard approaches, such as stable-isotope or fatty-acid analyses, can be used to infer the trophic position of predators in the food web, as well as potential switches in feeding sites (Elsdon, 2010; Hobson and Clark, 1992; Logan et al., 2006; MacNeil et al., 2005; Phillips and Eldridge, 2006; Taipale et al., 2011). Although they provide valuable information about trophic interactions, these methods do not reach a fine-scale resolution, usually lacking genus or species-level identification, which may be critical for the planning of conservation management actions (Bocher et al., 2000; Cherel et al., 2000; Deagle et al., 2007; Guest et al., 2009; Guillerault et al., 2017). In the last decade, parallel to the development and optimization of genomic techniques, DNA metabarcoding approaches using faecal material as a source of dietary DNA (de Sousa et al., 2019) have allowed the accurate identification of prey species within the diet of a wide variety of taxa including invertebrates (Kerley et al., 2018; Mollot et al., 2014; Pinol et al., 2014; Valentini et al., 2016) and vertebrates (Andriollo et al., 2019; Guillerault et al., 2017; Kamenova et al., 2018; Leray et al., 2015; Sullins et al., 2018).

The Westland petrel (*Procellaria westlandica*) is endemic to New Zealand and listed as an endangered species on the IUCN red list (BirdLife International, 2020). It is one of the few burrowing birds breeding on the main islands of New Zealand. This iconic species was once widespread in New Zealand (Waugh and Wilson, 2017; Wood and Otley, 2013), but its breeding distribution is now restricted to the West Coast of the South Island, within the Paparoa National Park and its surroundings (Jackson, 1958; Waugh and Wilson, 2017). Between May and June, females lay a single egg, which is incubated by both parents during 69 days (Warham, 1990). Chick-rearing is also carried out by both parents between September and November. After the breeding season, Westland petrels travel to South American waters (Baker and Coleman, 1977), where they remain until late March (March to November) (Landers et al., 2011). Regarding their foraging behaviour, Westland petrels are known to be nocturnal, but they occasionally feed during daytime (Waugh et al., 2018). Previous studies based on morphological analysis of regurgitates found that their most abundant prey items were fish, followed by cephalopods and crustaceans (Freeman, 1998; Imber, 1976). The diet of Westland petrels is therefore closely linked to fishing activity in New Zealand waters. However, it remains unclear whether fishing has a net positive or negative impact on *P. westlandica*. The overall population has increased significantly since the 70’s (Wood and Davis, 2003; Wood and Otley, 2013), together with the rise of fishing activity, potentially because of increase feeding on bycatch and other fishing waste. However, being trapped and killed in fishing nets is one of the main threats for *P. westlandica*, together with mammal predation, degradation of habitat and erosion of their nesting grounds (Taylor, 2000; Waugh et al., 2008; Waugh and Wilson, 2017).

The precise composition of the current diet of the Westland petrel is unknown, and potential temporal variations in diet throughout the breeding season have never been investigated. In this work, we present the first attempt to characterize the diet of this seabird through a DNA-based approach. To do this, we used a non-invasive DNA sampling approach (Lefort et al., 2019) by collecting faecal samples, and carrying out a DNA metabarcoding analysis using the 16S gene to identify prey items within the diet of the Westland petrel. This amplicon was chosen for the study as it has shown to be effective for the characterization of seabirds’ diet (Komura et al., 2018; McInnes et al., 2017; Young et al., 2020). The birds’ diet was compared between two breeding sub-colonies (1.5 km apart), and two different times (10 weeks apart), as another of our objectives was to describe potential differences between seasons and sub-colonies. Our hypothesis was that there would be differences in diet between early breeding season (before hatching) and late breeding season (after hatching or chick rearing), which would be consistent with switches in feeding and foraging behaviour. However, we did not expect to find significant differences in the diet of the different sub-colonies owing to their relatively close proximity. Our study also aimed at better understanding the impact that fishing activities have on Westland petrels by more accurately describing the composition of their diet.

## Methods

### Study area and sample collection

A total of 99 faecal samples were collected from two different sampling sites located in the West Coast of the South Island of New Zealand, the Paparoa National Park (NP) (-42.146317, 171.340293) (49 samples) and a private land (PL) (-42.164358, 171.337603) (50 samples) (Table S1). The collected samples were fresh and usually line-shaped, which could only correspond to faeces produced by birds as they landed on the previous day. Hence, each bird could only produce one of these faeces, and samples were considered independent. Very few older faecal samples were observed on the sites, as these were probably rapidly washed away in this extremely rainy location.

Forty-eight samples were collected before hatching (BH) on the 9^th^ and 10^th^ of July 2015, and 51 samples were collected during chick-rearing (CR) on the 22^nd^ and 23^rd^ of September 2015 (Table 1). To avoid cross-contamination, each fresh faecal sample was collected using an individual sterile cotton swab and placed in a clean, single-use Ziplock bag. Samples were then placed in a cooled icebox for transportation to the laboratory (within the following two days), where they were stored at -80°C until DNA extraction. Leaf litter samples were also collected to serve as negative controls.

**Table 1.** OTU list after filtering the contaminants, low quality sequences and the sequences that gave no hits. For each OTU, the taxonomical classification is given, together with the standard parameters provided by the BLAST search against the NCBI database. The penultimate column indicates whether the OTU was identified in previous studies or not. The last column gives the depth at which each OTU is naturally found, and coloured rows indicates OTUs whose depth range overlaps with the dive depth of the Westland petrel.

| OTU_ID | Phylum | Class | Order | Family/Superfamily | Genus | Species | Size | E-value | % of identity | Alignment length | Query cover | Previously identified | Depth (m) |
| --- | --- | --- | --- | --- | --- | --- | --- | --- | --- | --- | --- | --- | --- |
| OTU_43 | Arthropoda | Branchiopoda | Anostraca | Artemiidae | *Artemia* | *Artemia franciscana* | 2482 | 2.36E-71 | 1 | 151 | 0.9934 | NO | 0.1-0.6 |
| OTU_81 | Arthropoda | Maxillopoda | Calanoida | Candaciidae | *Candacia* | *Candacia armata* | 761 | 2.13E-50 | 1 | 113 | 0.9912 | NO | - |
| OTU_66 | Arthropoda | Malacostraca | Decapoda | Penaeidae | *Penaeus* | *Penaeus merguensis* | 1216 | 7.47E-82 | 1 | 170 | 0.9941 | NO | 10-45 |
| OTU_65 | Arthropoda | Malacostraca | Decapoda | Pilumnidae | *Pilumnus* | *Pilumnus hirtellus* | 1269 | 2.57E-76 | 0.9939 | 163 | 0.9939 | NO | 10-80 |
| OTU_7 | Arthropoda | Malacostraca | Amphipoda | Talitroidea | not identified | not identified | 92506 | 8.65E-16 | 0.8485 | 99 | 0.6528 | NO | 0-0.1 |
| OTU_16 | Arthropoda | Malacostraca | Amphipoda | Talitroidea | not identified | not identified | 50055 | 1.92E-07 | 0.8000 | 95 | 0.6370 | NO | 0-0.1 |
| OTU_17 | Arthropoda | Malacostraca | Amphipoda | Talitroidea | not identified | not identified | 28822 | 6.68E-17 | 0.8092 | 131 | 0.8819 | NO | 0-0.1 |
| OTU_28 | Arthropoda | Malacostraca | Amphipoda | Talitroidea | not identified | not identified | 11687 | 3.99E-19 | 0.8014 | 146 | 0.9931 | NO | 0-0.1 |
| OTU_212 | Arthropoda | Malacostraca | Amphipoda | Talitroidea | not identified | not identified | 75 | 1.92E-07 | 0.8061 | 98 | 0.6370 | NO | 0-0.1 |
| OTU_229 | Arthropoda | Malacostraca | Amphipoda | Talitroidea | not identified | not identified | 14954 | 8.93E-06 | 0.7895 | 95 | 0.6370 | NO | 0-0.1 |
| OTU_304 | Arthropoda | Malacostraca | Amphipoda | Talitroidea | not identified | not identified | 46 | 3.05E-10 | 0.7813 | 128 | 0.8936 | NO | 0-0.1 |
| OTU_309 | Arthropoda | Malacostraca | Amphipoda | Talitroidea | not identified | not identified | 44 | 3.97E-24 | 0.8219 | 146 | 0.9931 | NO | 0-0.1 |
| OTU_323 | Arthropoda | Malacostraca | Amphipoda | Talitroidea | not identified | not identified | 192 | 6.64E-22 | 0.8321 | 131 | 0.8819 | NO | 0-0.1 |
| OTU_402 | Arthropoda | Malacostraca | Amphipoda | Talitroidea | not identified | not identified | 23 | 1.92E-07 | 0.8081 | 99 | 0.6370 | NO | 0-0.1 |
| OTU_405 | Arthropoda | Malacostraca | Amphipoda | Talitroidea | not identified | not identified | 138 | 1.82E-27 | 0.8356 | 146 | 0.9930 | NO | 0-0.1 |
| OTU_406 | Arthropoda | Malacostraca | Amphipoda | Talitroidea | not identified | not identified | 28 | 1.92E-07 | 0.8061 | 98 | 0.6370 | NO | 0-0.1 |
| OTU_416 | Arthropoda | Malacostraca | Amphipoda | Talitroidea | not identified | not identified | 777 | 8.65E-16 | 0.8557 | 97 | 0.6389 | NO | 0-0.1 |
| OTU_434 | Arthropoda | Malacostraca | Amphipoda | Talitroidea | not identified | not identified | 660 | 3.08E-15 | 0.8485 | 99 | 0.6503 | NO | 0-0.1 |
| OTU_471 | Arthropoda | Malacostraca | Amphipoda | Talitroidea | not identified | not identified | 23 | 1.92E-07 | 0.8000 | 95 | 0.6370 | NO | 0-0.1 |
| OTU_524 | Arthropoda | Malacostraca | Amphipoda | Talitroidea | not identified | not identified | 21 | 2.41E-21 | 0.8182 | 143 | 0.9724 | NO | 0-0.1 |
| OTU_529 | Arthropoda | Malacostraca | Amphipoda | Talitroidea | not identified | not identified | 23 | 1.92E-07 | 0.8000 | 95 | 0.6370 | NO | 0-0.1 |
| OTU_557 | Arthropoda | Malacostraca | Amphipoda | Talitroidea | not identified | not identified | 148 | 8.93E-06 | 0.7895 | 95 | 0.6370 | NO | 0-0.1 |
| OTU_593 | Arthropoda | Malacostraca | Amphipoda | Talitroidea | not identified | not identified | 16 | 1.86E-17 | 0.8043 | 138 | 0.9306 | NO | 0-0.1 |
| OTU_615 | Arthropoda | Malacostraca | Amphipoda | Talitroidea | not identified | not identified | 146 | 3.99E-19 | 0.8041 | 148 | 0.9931 | NO | 0-0.1 |
| OTU_636 | Arthropoda | Malacostraca | Amphipoda | Talitroidea | not identified | not identified | 47 | 1.92E-07 | 0.8000 | 95 | 0.6370 | NO | 0-0.1 |
| OTU_662 | Arthropoda | Malacostraca | Amphipoda | Talitroidea | not identified | not identified | 312 | 5.12E-18 | 0.8106 | 132 | 0.8951 | NO | 0-0.1 |
| OTU_666 | Arthropoda | Malacostraca | Amphipoda | Talitroidea | not identified | not identified | 646 | 5.08E-23 | 0.8333 | 132 | 0.8951 | NO | 0-0.1 |
| OTU_681 | Arthropoda | Malacostraca | Amphipoda | Talitroidea | not identified | not identified | 32 | 1.4337E-13 | 0.8404 | 94 | 0.6294 | NO | 0-0.1 |
| OTU_684 | Arthropoda | Malacostraca | Amphipoda | Talitroidea | not identified | not identified | 13 | 3.14E-15 | 0.7970 | 133 | 0.9034 | NO | 0-0.1 |
| OTU_691 | Arthropoda | Malacostraca | Amphipoda | Talitroidea | not identified | not identified | 64 | 1.11E-19 | 0.8095 | 147 | 0.9931 | NO | 0-0.1 |
| OTU_724 | Arthropoda | Malacostraca | Amphipoda | Talitroidea | not identified | not identified | 3816 | 0.00011547 | 0.7849 | 93 | 0.6233 | NO | 0-0.1 |
| OTU_785 | Arthropoda | Malacostraca | Amphipoda | Talitroidea | not identified | not identified | 67 | 8.51E-21 | 0.8095 | 147 | 0.9930 | NO | 0-0.1 |
| OTU_786 | Arthropoda | Malacostraca | Amphipoda | Talitroidea | not identified | not identified | 237 | 5.16E-13 | 0.7879 | 132 | 0.8951 | NO | 0-0.1 |
| OTU_788 | Arthropoda | Malacostraca | Amphipoda | Talitroidea | not identified | not identified | 25 | 4.93E-18 | 0.8071 | 140 | 0.9712 | NO | 0-0.1 |
| OTU_802 | Arthropoda | Malacostraca | Amphipoda | Talitroidea | not identified | not identified | 15 | 1.86E-17 | 0.8586 | 99 | 0.6528 | NO | 0-0.1 |
| OTU_811 | Arthropoda | Malacostraca | Amphipoda | Talitroidea | not identified | not identified | 8 | 1.84E-17 | 0.7959 | 147 | 0.9930 | NO | 0-0.1 |
| OTU_828 | Arthropoda | Malacostraca | Amphipoda | Talitroidea | not identified | not identified | 10 | 3.99E-19 | 0.8027 | 147 | 0.9931 | NO | 0-0.1 |
| OTU_869 | Arthropoda | Malacostraca | Amphipoda | Talitroidea | not identified | not identified | 67 | 2.40E-11 | 0.8333 | 90 | 0.6084 | NO | 0-0.1 |
| OTU_894 | Arthropoda | Malacostraca | Amphipoda | Talitroidea | not identified | not identified | 8 | 1.92E-07 | 0.8081 | 99 | 0.6370 | NO | 0-0.1 |
| OTU_908 | Arthropoda | Malacostraca | Amphipoda | Talitroidea | not identified | not identified | 1592 | 3.99E-19 | 0.8014 | 146 | 0.9931 | NO | 0-0.1 |
| OTU_914 | Arthropoda | Malacostraca | Amphipoda | Talitroidea | not identified | not identified | 83 | 3.01E-05 | 0.7957 | 93 | 0.6187 | NO | 0-0.1 |
| OTU_936 | Arthropoda | Malacostraca | Amphipoda | Talitroidea | not identified | not identified | 7 | 3.50E-15 | 0.8030 | 132 | 0.7975 | NO | 0-0.1 |
| OTU_937 | Arthropoda | Malacostraca | Amphipoda | Talitroidea | not identified | not identified | 6 | 2.37E-10 | 0.8533 | 75 | 0.6239 | NO | 0-0.1 |
| OTU_949 | Arthropoda | Malacostraca | Amphipoda | Talitroidea | not identified | not identified | 5 | 1.41E-23 | 0.8900 | 100 | 0.6713 | NO | 0-0.1 |
| OTU_979 | Arthropoda | Malacostraca | Amphipoda | Talitroidea | not identified | not identified | 20 | 8.79E-11 | 0.7762 | 143 | 0.9517 | NO | 0-0.1 |
| OTU_1000 | Arthropoda | Malacostraca | Amphipoda | Talitroidea | not identified | not identified | 8 | 2.38E-16 | 0.8485 | 99 | 0.6643 | NO | 0-0.1 |
| OTU_1055 | Arthropoda | Malacostraca | Amphipoda | Talitroidea | not identified | not identified | 56 | 4.02E-14 | 0.8523 | 88 | 0.6042 | NO | 0-0.1 |
| OTU_1057 | Arthropoda | Malacostraca | Amphipoda | Talitroidea | not identified | not identified | 5 | 2.77E-15 | 0.8049 | 123 | 0.9167 | NO | 0-0.1 |
| OTU_1059 | Arthropoda | Malacostraca | Amphipoda | Talitroidea | not identified | not identified | 68 | 4.93E-18 | 0.8082 | 146 | 0.9928 | NO | 0-0.1 |
| OTU_1065 | Arthropoda | Malacostraca | Amphipoda | Talitroidea | not identified | not identified | 206 | 1.48E-08 | 0.7963 | 108 | 0.7123 | NO | 0-0.1 |
| OTU_1104 | Arthropoda | Malacostraca | Amphipoda | Talitroidea | not identified | not identified | 6 | 6.32E-17 | 0.8721 | 86 | 0.6159 | NO | 0-0.1 |
| OTU_1144 | Arthropoda | Malacostraca | Amphipoda | Talitroidea | not identified | not identified | 28 | 3.96E-19 | 0.8014 | 146 | 0.9930 | NO | 0-0.1 |
| OTU_59 | Chordata | Actinopterygii | Anguilliformes | Nettastomatidae | not identified | not identified | 1636 | 6.92E-64 | 0.8595 | 242 | 0.9958 | NO | Deep-sea |
| OTU_169 | Chordata | Actinopterygii | Gadiformes | Euclichthyidae | *Euclichthys* | *Euclichthys polynemus* | 148 | 2.04E-103 | 0.9953 | 212 | 0.9953 | NO | 250-920 |
| OTU_22 | Chordata | Actinopterygii | Gadiformes | Macrouridae | *Lepidorhynchus* | *Lepidorhynchus denticulatus* | 17801 | 7.50E-108 | 1 | 217 | 0.9954 | Freeman, 1998; Imber, 1976 | 270-450 |
| OTU_54 | Chordata | Actinopterygii | Gadiformes | Macrouridae | *Coelorinchus* | *Coelorinchus oliverianus* | 2067 | 9.59E-107 | 1 | 215 | 0.9953 | Freeman, 1998 (genus level) | 400-600 |
| OTU_216 | Chordata | Actinopterygii | Gadiformes | Macrouridae | *Coelorinchus* | *Coelorinchus fasciatus* | 107 | 2.08E-103 | 0.9907 | 215 | 0.9953 | Freeman, 1998 (genus level); Imber, 1976 (genus level) | 400-600 |
| OTU_1075 | Chordata | Actinopterygii | Gadiformes | Macrouridae | *Coelorinchus* | *Coelorinchus oliverianus* | 12 | 1.27E-95 | 0.9721 | 215 | 0.9861 | Freeman, 1998 (family level) | 400-600 |
| OTU_12 | Chordata | Actinopterygii | Gadiformes | Merlucciidae | *Macruronus* | *Macruronus novaezelandiae* | 49195 | 9.59E-107 | 1 | 215 | 0.9953 | Freeman, 1998 | 200-700 |
| OTU_14 | Chordata | Actinopterygii | Gadiformes | Merlucciidae | *Merluccius* | *Merluccius australis* | 46909 | 3.43E-106 | 1 | 214 | 0.9953 | Freeman, 1998 (family level) | 28-1000 |
| OTU_808 | Chordata | Actinopterygii | Gadiformes | Merlucciidae | *Macruronus* | *Macruronus* sp. | 66 | 2.55E-77 | 0.9302 | 215 | 0.9950 | Freeman, 1998 | 0-1000 |
| OTU_1008 | Chordata | Actinopterygii | Gadiformes | Merlucciidae | *Merluccius* | *Merluccius productus* | 14 | 1.32E-27 | 0.9867 | 75 | 0.6549 | NO | 0-1000 |
| OTU_1086 | Chordata | Actinopterygii | Gadiformes | Merlucciidae | not identified | not identified | 44 | 3.61E-66 | 0.8884 | 215 | 0.9953 | Freeman, 1998 (family level) | 28-1000 |
| OTU_107 | Chordata | Actinopterygii | Gadiformes | Moridae | *Mora* | *Mora moro* | 466 | 3.43E-106 | 1 | 214 | 0.9953 | Freeman, 1998 (family level) | 450-2500 |
| OTU_38 | Chordata | Actinopterygii | Ophidiiformes | Ophidiidae | *Genypterus* | *Genypterus blacodes* | 5711 | 2.68E-107 | 1 | 216 | 0.9954 | NO | 300-550 |
| OTU_962 | Chordata | Actinopterygii | Ophidiiformes | Ophidiidae | *Genypterus* | *Genypterus* sp. | 10 | 4.55E-95 | 0.9676 | 216 | 0.9907 | NO | 22-1000 |
| OTU_15 | Chordata | Actinopterygii | Perciformes | Triglidae | *Lepidotrigla* | *Lepidotrigla modesta* | 45382 | 4.46E-105 | 0.9953 | 215 | 0.9953 | NO | 10-300 |
| OTU_96 | Chordata | Actinopterygii | Scombriformes | Gempyidae | *Rexea* | *Rexea* sp. | 565 | 2.15E-93 | 0.9589 | 219 | 0.9954 | NO | 100-800 |
| OTU_18 | Chordata | Actinopterygii | Scombriformes | Trichiuridae | *Trichiuridae environmental sample* | not identified | 26490 | 1.74E-94 | 0.9515 | 227 | 0.9956 | Imber, 1976 (family level) | 0-1600 |
| OTU_163 | Chordata | Actinopterygii | Zeiformes | Cyttidae | *Cyttus* | *Cyttus traversi* | 189 | 9.59E-107 | 1 | 215 | 0.9953 | Freeman, 1998 (genus level) | 200-978 |
| OTU_86 | Chordata | Actinopterygii | Zeiformes | Zenionidae | *Capromimus* | *Capromimus abbreviatus* | 629 | 9.59E-107 | 1 | 215 | 0.9953 | NO | 87-500 |
| OTU_19 | Mollusca | Cephalopoda | Octopoda | Octopodidae | *Octopus* | *Octopus vulgaris* | 20934 | 1.55E-78 | 1 | 164 | 0.9939 | Freeman, 1998 (genus level) | 0-100 |
| OTU_1019 | Mollusca | Cephalopoda | Octopoda | Octopodidae | *Octopus* | *Octopus* sp. | 24 | 1.21E-64 | 0.9515 | 165 | 0.9939 | Freeman, 1998 (genus level) | 0-100 |
| OTU_8 | Mollusca | Cephalopoda | Teuthida | Loliginidae | not identified | not identified | 104188 | 7.91E-87 | 1 | 179 | 0.9944 | NO | 0-400 |
| OTU_60 | Mollusca | Cephalopoda | Teuthida | Loliginidae | not identified | not identified | 2092 | 2.82E-86 | 1 | 178 | 0.9944 | NO | 0-400 |
| OTU_1003 | Mollusca | Cephalopoda | Teuthida | Loliginidae | not identified | not identified | 4 | 3.81E-60 | 0.9121 | 182 | 0.9944 | NO | 0-400 |
| OTU_1040 | Mollusca | Cephalopoda | Teuthida | Loliginidae | not identified | not identified | 8 | 2.88E-66 | 0.9375 | 176 | 0.9831 | NO | 0-400 |
| OTU_1125 | Mollusca | Cephalopoda | Teuthida | Loliginidae | not identified | not identified | 114 | 7.52E-77 | 1 | 161 | 0.9412 | NO | 0-400 |
| OTU_80 | Mollusca | Cephalopoda | Oegopsida | Histioteuthidae | not identified | not identified | 660 | 6.68E-42 | 0.8919 | 148 | 0.9797 | Freeman, 1998; Imber, 1976 | 300-400 |

### DNA extraction, PCR amplification and sequencing

For each faecal sample, we performed a DNA extraction on one small subsample collected with a cotton swab. We used the QIAamp DNA Stool Mini Kit (Trevelline et al., 2018, 2016) for which we followed the manufacturer’s protocol (Handbook from 03/2014, reference: 1081060_HB_LS) with few modifications. In brief, half volumes of all reagents were used and the extraction was carried out in 1.5 ml tubes, instead of 2 ml tubes. In addition, after adding half an InhibitEx Tablet, we performed only one centrifugation, rather than 2 (Steps 6 and 7 in the protocol were joined). Later, on step 13, we mixed 200 µl of ethanol by pipetting and 600 µl of the mix were added to the column. From step 14, volumes recommended by the manufacturer’s protocol were used. Finally, samples were eluted in 100 µl of elution buffer (AE) and DNA extracts stored at -20°C.

Later, two different PCR amplifications were performed from each DNA extract. First, we used a pair of primers specific for Chordata (fish) (16S1F, 16S2R), which amplifies 155 bp of the 16S gene (Deagle et al., 2009, 2005). Second, we used a pair of primers originally designed for Malacostraca (crustaceans) (Chord_16S_F_TagA, Chord_16S_R_Short) (Deagle et al., 2009) but also known to amplify cephalopods DNA in an efficient manner (Olmos-Pérez et al., 2017). This second pair of primers targets a 205 bp region of the 16S gene (Deagle et al., 2009). These two pairs of primers were tagged with sequence fragments, which are complementary to the illumina ligation adaptor. These primers were chosen to allow the detection of a wide range of potential prey, including the main taxa identified morphologically in previous studies, namely fish, cephalopods and crustaceans (Freeman, 1998; Imber, 1976). PCR conditions for both primer pairs were the same as in Olmos-Perez et al. (2017), with the exception of the Taq polymerase. Here, the FirePOLE® Taq polymerase was used for all amplifications, following manufacturer’s protocol (Solis BioDyne). Negative controls containing DNA-free water were added to each PCR run. After checking the results in a 1.5% agarose gel, PCR products were purified using AMPURE magnetic beads, following the manufacturer’s standard protocol (1.8 μL AMPure XP per 1.0 μL of sample) and, finally, for each sample, both PCR products were equalized at 2 ng/µL and pooled. Second stage PCR amplifications and subsequent sequencing steps were carried out by New Zealand Genomics Limited (NZGL, University of Otago) according to the Illumina “16S Metagenomic Sequencing Library Preparation Manual” Rev B. The resulting amplicons indexed with Nextera adapters (Index Set C) in unique combinations were arranged in two plates.

Sequencing was performed by NZGL on Illumina MiSeq 2x300bp reads (Illumina MiSeq v3 8 reagent kit).

### Bioinformatic library filtering

Metabarcoding library filtering was performed using a toolbox of software. First, we trimmed the two pairs of primers separately using *cutadapt* (Martin, 2011), leaving the maximum error rate as default (e=0.1). At this point, we had two sets of trimmed sequences. The following filtering steps were done twice in each pair of primers. Pair reads were merged using *PEAR* (Zhang et al., 2014), setting the minimum quality threshold (*Phred* score) at 30 (-q 30). After merging, all sequences were merged into a single *fastq* file using the *sed* command and all the subsequent steps were performed using *vsearch v2.8.1* (Rognes et al., 2016).

This step was followed by the quality rate filtering, which aims to remove sequences with sequencing errors, using the *fastx_filter* command (*fastq_maxee*=1). The library was dereplicated using the *derep_fulllength* command and, after, frequency errors were detected and deleted using again the *fastx_filter* command (*minsize*=2) in order to delete the singletons, as such low frequency variants are likely to be PCR errors. The following step was filtering sequences by length (indel filtering) with the command *fastx_filter* (*fastq_minlen*=50, *fastq_maxlen*=325). At this stage, merged sequences that were shorter than 50bp or longer that 325bp were discarded. This step was followed by the filtering *de novo* of potential chimeras using the *uchime_denovo* command. After this step, we obtained a *fasta* file, with Amplicon Single Variants (ASVs). Next, we performed the Operational Taxonomic Unit (OTU) clustering, applying the centroid-based greedy clustering algorithm with a cut-off threshold of 97% (Xiong and Zhan, 2018) using the *cluster_size* command (*id* 0.97), and obtained a *fasta* file with all the OTUs present in the sampling. Finally, we mapped the reads in each sample to OTUs in order to obtain an OTU table, using *search_exact* command. Thus, at this point of the pipeline, we obtained two output files, an OTU table and a *fasta* file with the subsequent sequence of all the OTU sequences.

All OTUs were compared to the NCBI database using NCBI BLAST web interface (Johnson et al., 2008) and the pertinent multiple-file JSON was downloaded from this web interface. We then used a customized R script, based on the functions *fromJSON* and *classification* from R packages *rjson* (Couture-Beil and Couture-Beil, 2018) and *taxize* (Chamberlain and Szöcs, 2013), respectively, to retrieve the best hit from the taxonomic classification of each clustered OTU from the NCBI database. Moreover, we performed SINTAX classification against the MIDORI database (Leray et al., 2018). For that purpose, we used the *vsearch* commands *makeudb_search* to convert the database, which was downloaded from the MIDORI website in *fasta* file, into database format and *sintax* to retrieve the taxonomic classification.

Regarding the taxonomic assignment, we discarded OTUs with BLAST query coverage under 60% or BLAST identities lower than 75%. The number of reads present in the negative control were subtracted from each sample as they were considered as potential contaminations. Also, singletons among samples and OTUS were also considered as potential contamination or artifacts, and removed from the dataset as were any OTUs matching to Westland Petrel (Brown et al., 2015; Gobet et al., 2010; Lindahl et al., 2013; Majaneva et al., 2015; Shade et al., 2012). We also filtered the taxonomic assignment table, discarding every OTU which was classified as prokaryotes, fungi, insects, mammals and the Westland petrel itself, as they could not be a potential prey for biological reasons. Potential prey OTUs within the phyla Arthropoda, Chordata and the Mollusca families Octopodidae and Histiotheutidae were assigned using the following criteria to taxonomical categories: OTUs with identity higher than 97% were determined at species level, OTUs between 93 and 97% were assigned to genus level, and OTUs with identity below 93% were assigned to family level. In the case of the Mollusca family Loliginidae, we obtained a taxonomic assignment corresponding to species not present in New Zealand’s waters. Therefore, OTUs were aligned with 100 Loliginidae sequences retrieved from GenBank (Benson et al., 2012). This alignment revealed that the 16S fragment was exactly the same for several genus of this family, meaning that this amplicon fragment does not have sufficient resolution to resolve genus and species identity within this family. Thus, OTUs matching the Loliginidae family were only assigned to family level, regardless of the percentage of identity retrieved from the BLASTn taxonomic assignment.

### Biodiversity analyses

In order to evaluate the impact of commercial marine species on the diet of *P. westlandica*, we collected ecological information from FishBase (Froese and Pauly, 2010) and SeaLifeBase (Palomares and Pauly, 2010) to determine the distribution of each prey taxa. Considering that *P. westlandica* is able to dive up to 15 m for fishing (Waugh et al., 2018), we specifically looked for information about the depth at which the prey species are usually present (shallow versus deep sea) and whether they were naturally reachable for the Westland petrel. We also checked whether those prey species had been detected in previous publications (Table 1). To measure the completeness of our sampling, we evaluated the total richness of prey in the diet of *P. westlandica*, using a rarefaction curve and a bootstrap estimator with the function *specpool* in the R package *vegan* (Oksanen et al., 2013). Moreover, as a measure of the quality of our sequencing, we plotted the number of sequence reads per OTU detected (Fig. S3) and the cumulative frequency of OTU detected in relation to the number of sequence reads produced (Fig. S4). The diet of Westland petrels was described using two different metrics. First, we calculated the Frequency of Occurrence (FOO), which gives the information about the number of samples in which an OTU is present. This was calculated by transforming the number of reads to presence/absence (1-0) and, subsequently, summing the counts of samples that contain a given prey item (OTU), expressed as a proportion of all samples (Deagle et al., 2019). Second, we calculated the Relative Read Abundance (RRA), which is the proportion of reads obtained for each prey item (OTU) in each sample (Deagle et al., 2019), which was calculated using the OTU table of abundances. Both metrics were computed with customized scripts using the R package *dplyr* (Wickham et al., 2021). FOO and RRA were calculated overall to describe the diet of Westland petrels as a species, and also compared between seasons: before hatching (BH) versus chick-rearing (CR); and between sub-colonies: natural park (NP) versus private land (PL).

To estimate the effects of seasonality and sub-colony location on diet diversity and composition, we computed a negative binomial Generalized Linear Model (GLM) (McCullagh and Nelder, 1989) with a log link function, applying the function *manyglm* from the R package *mvabund* (Wang et al., 2017). Two different GLM analyses were performed, one with read abundance as the dependant variable and one with occurrences as the dependant variable. For both GLM analyses, the predictor variables were season (two factor levels: BH and CR) and site (two factor levels: NP and PL) as well as the interaction between these variables. An analysis of Deviance (Dev) was performed to test the fitness of the model, with 999 bootstraps iterations as a resampling method (Davison and Hinkley, 1997), using the function *anova.manyglm* from the package *mvabund* (Wang et al., 2017). Moreover, an ordination to visualize the differences in community composition between the two seasons (BH and CR) and the two sub-colonies or sites (NP and PL) was computed and plotted using the *cord* function from the R package *ecoCopula* (Popovic et al, 2021).

Finally, we estimated and plotted the standard alpha diversity, as a proxy for prey species richness, comparing the two factors studied, season and site. For that purpose, we used the functions *diversity* and *plot_richness* from R packages *vegan* and *phyloseq* (McMurdie and Holmes, 2012; Oksanen et al., 2013), respectively. In addition, we computed pairwise comparisons between the alpha diversity values (Simpson) of the group levels through the pairwise Wilcoxon rank sum test (Gehan, 1965), using the function *pairwise.wilcox.text* from the R package *stats* (Team and others, 2013).

## Results

### Amplification success and library quality

All 98 samples were successfully amplified with both pairs of primers and sequenced with Illumina MiSeq. We obtained a total of 9,847,628 raw reads. After trimming with *cutadapt* the Malacostraca pair of primers (Deagle et al., 2005), we obtained 7,085,188 reads and 3,010,097 merged reads (84,97% of the raw data was merged). In the next step, we obtained 3,010,097 quality filtered reads, which resulted in 150,973 dereplicated unique sequences. Finally, after indel and chimera filtering, we obtained 31,691 ASVs, which were clustered in 1,147 OTUs (Fig.S1). In the case of the Chordata pair of primers, after trimming we obtained 321,240 reads which resulted in 20 OTUs, in which only 1 OTU with 3 reads was different from the OTU set obtained with the Malacostraca pair of primers. Thus, the sequences from the Chordata primers were discarded, and only the reads obtained with the Malacostraca pair of primers were retained for subsequent analyses. The 1,147 OTUs comprised 2,567,254 reads, from which 127,088 (243 OTUs) were considered as contaminants (not potential prey) 560,586 reads (102 OTUs) were considered as low-quality assignment (query cover < 60% and percentage of identity < 75%), 1,371,994 reads (723 OTUs) did not match against GenBank, and 507,231 were considered as the reads of potential prey of the Westland petrels. These potential prey reads belonged to 79 OTUs (Table 1), and 17 samples had only unassigned or undetermined OTUs and, hence, were not used in the subsequent analyses. We were not able to recover any additional assignment from MIDORI compared to those obtained from GenBank. Thus, this information was discarded.

### Characterization of the diet of *P. westlandica*

Species richness estimation (based on a bootstrap analysis) suggested that our sampling captured 88.6% of the total diversity of prey items within the diet of *P. westlandica* (Fig.S2). The number of sequence reads per OTUs detected (Fig.S3) and the cumulative frequency of the OTUs detected (Fig.S4), which are both measures of sequencing depth, were sufficient to characterize the diet of the Westland petrel. Out of the 79 OTUs recovered by metabarcoding, 24.02% (19 OTUs, 195,358 reads) were identified to species level, 5.06% (4 OTUs, 29,089 reads) were identified to genus level and 70.89% (56 OTUs, 316,587 reads) were identified to family level (Table 1).

Arthropods (crustaceans in this case) were the most common prey in the diet of Westland petrels, being present in 62.03% of the samples (FOO) and represented 45.57% of the sequences (RRA) and 65.82% of the OTUs. Actinopterygii (bony fish) were next, being present in 59.49% of the samples and comprising 42.13% of all sequences and 24.05% of all OTUs. Finally, cephalopods were present in 53.16% of the samples and made up 12.29% of the sequences and 10.12% of the OTUs (Fig.1). Within arthropods, Talitridae (landhoppers and sandhoppers) were by far the most abundant taxa. Although there are marine talitrids (Fenwick, 2001; Lowry and Bopiah, 2012), there is insufficient information in the databases and possible faulty matches as amphipodan taxonomy is challenging and under continuous change. That is the reason why, in this study, we will use the higher-level taxonomic assignment until superfamily Talitroidea. They were present in 58.23% of the samples and made up 44.35% of the sequences. Other minor arthropod taxa were identified, such as the families Pilumnidae (pilumnid crabs) and Penaeidae (penaeid shrimps), among others (<1% total reads; Table 2). With the exception of four OTUs, which were identified to species level, arthropods were identified to family level.

**Figure 1.**
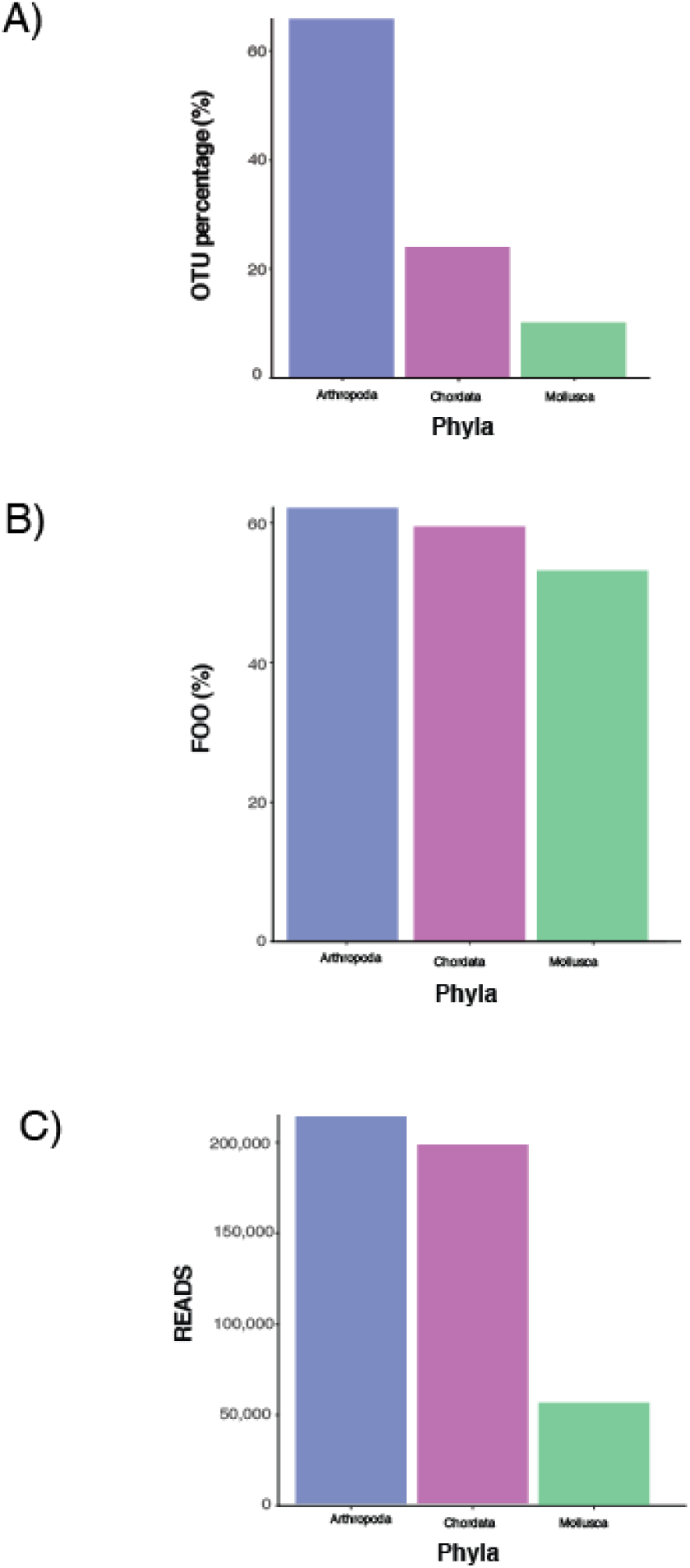
Prey phyla identified using three different biodiversity metrics: A) Number of OTUs as a proxy of diversity, B) Frequency of occurrence (FOO) refers to the percentage of samples in which the prey item is present and C) Read abundance.

**Table 2.**
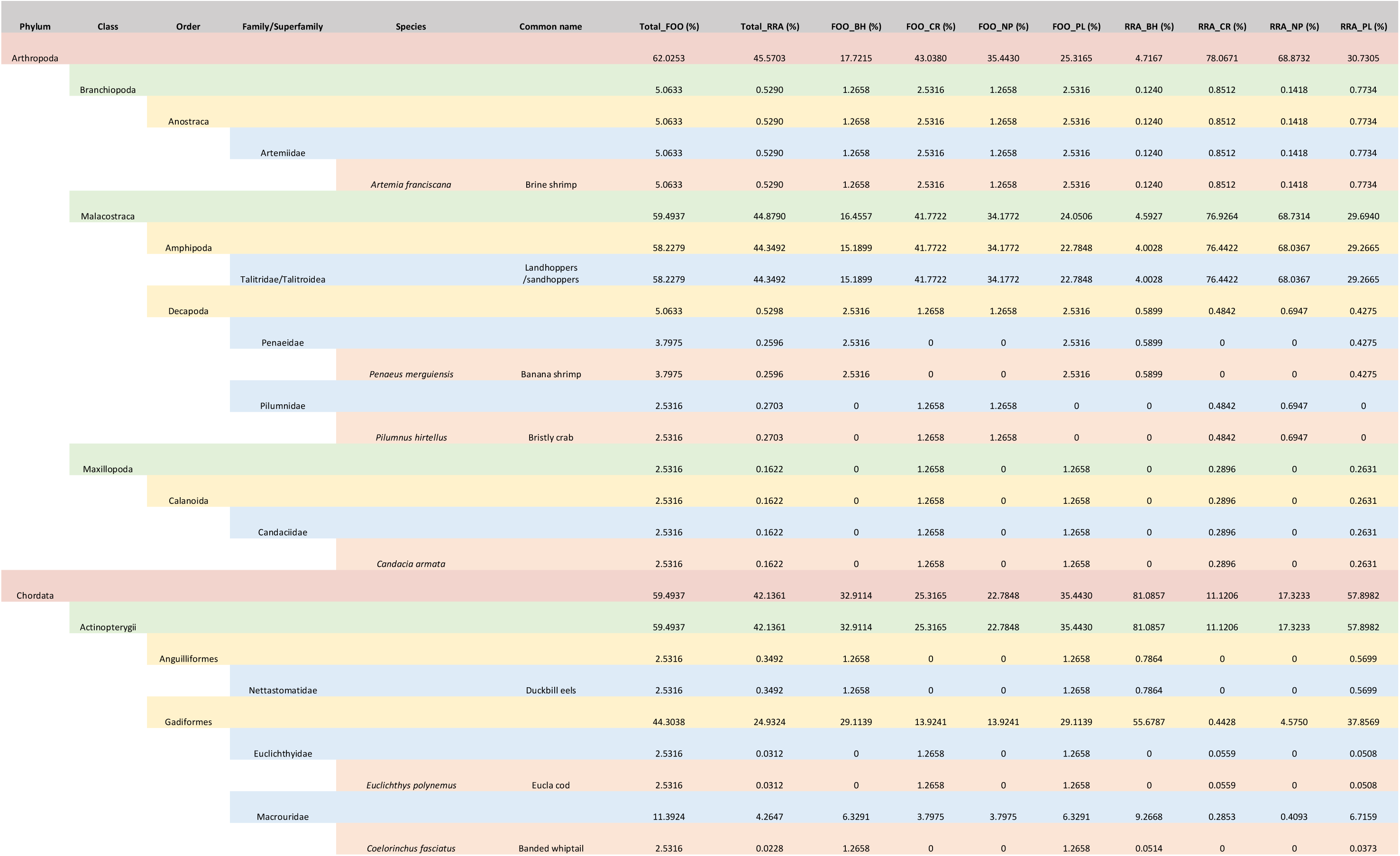
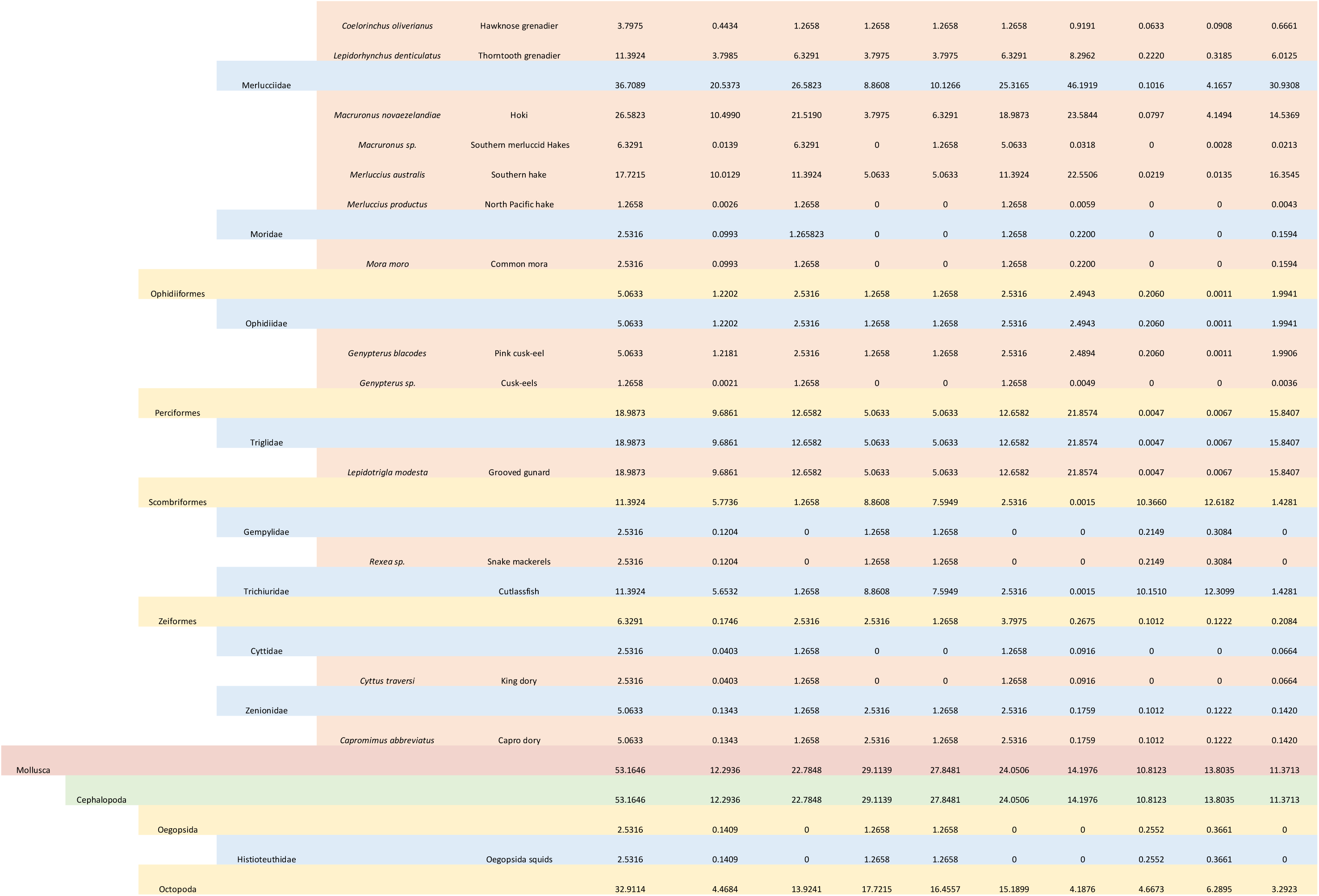
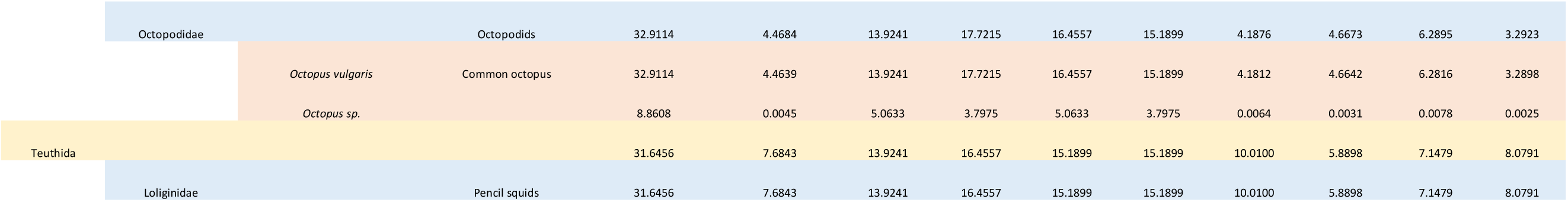
Taxonomical classification of the prey items of *P. westlandica* until family level with its corresponding Relative Read Abundance (RRA) and Frequency Of Occurrence (FOO) values for the whole sampling and showing the differences among: A) the two different seasons, Before Hatching (BH) and Chick-rearing (CR) and B) the two different sites, the Paparoa Natural Park (NP) and the Privat Land (PL) in the surroundings.

Within Chordata (ray-finned fish in this case), Hoki (*Macruronus novaezelandiae*) was the most common species as it was present in 26.58% of the samples and represented 10.5% of all sequences. The Cocky gurnard (*Lepidotrigla modesta*) and the Southern hake (*Merluccius australis*) were also important prey items, being present in 18.99% and 17.72 % of the samples and comprising 9.69% and 10.01% of all sequences, respectively. Next were cutlassfish, identified to family level (Trichiuridae), and the Thorntooth grenadier (*Lepidorhynchus denticulatus*) both present in 11.39% of the samples and comprising 5.77% and 3.8% of all sequences, respectively. As in the case of arthropods, we detected few other minor taxa, such as the Pink cusk-eel (*Genypterus blacodes*) or the Hawknose grenadier (*Coelorinchus oliverianus*), among others (around 1% of the reads: Table 2; Fig.2). Out of 19 OTUs of Actinopterygii, three OTUs were identified to genus level, two OTUs were identified to family and the remaining 13 OTUs were identified to species level (Table 1).

**Figure 2.**
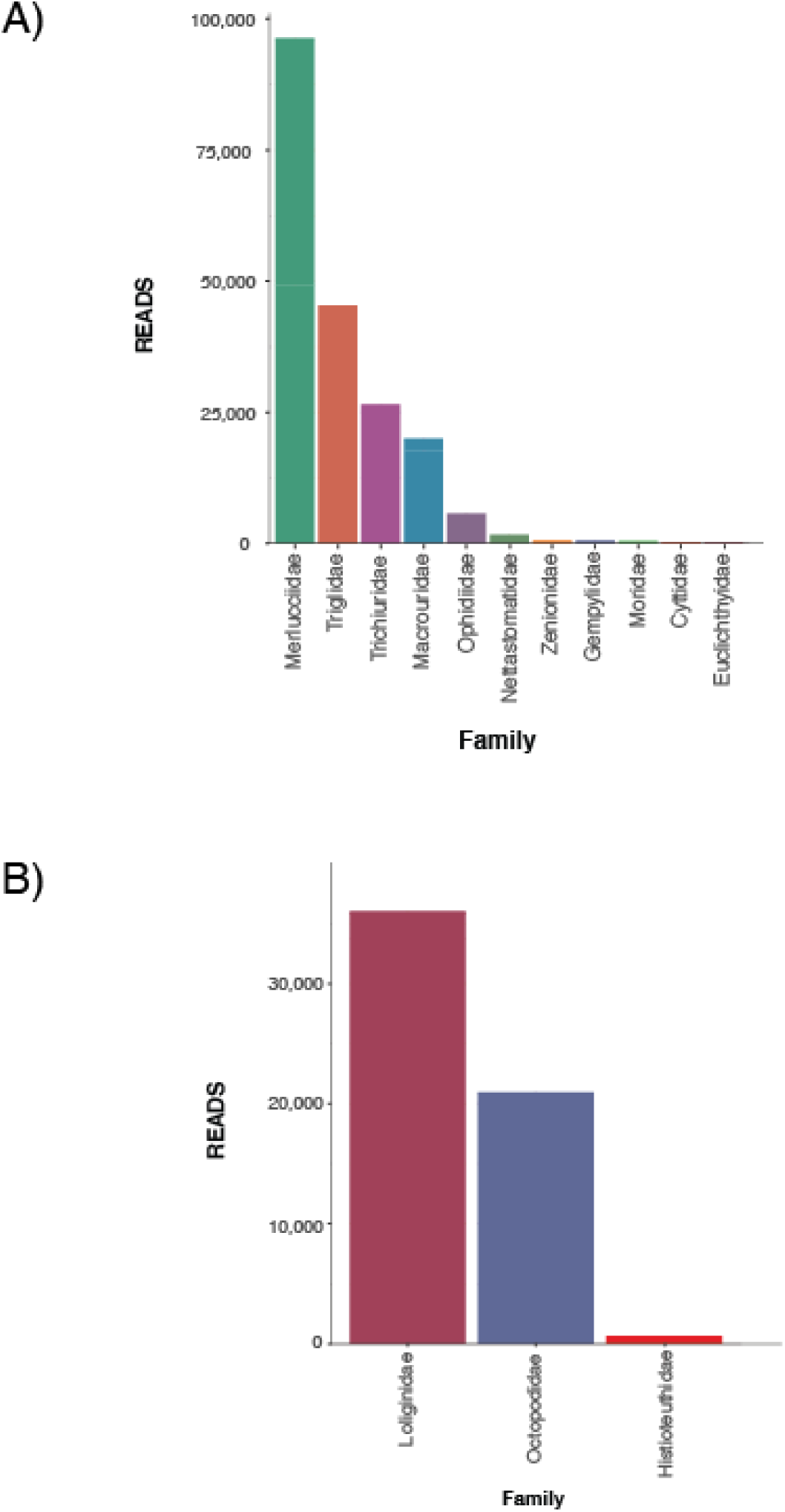
Read abundance classified by family for A) Fish prey items B) Cephalopod prey items

According to our results, within cephalopods, eight different OTUs were identified as prey items, six of which were assigned to family level, one to genus level and one to species level (Table 1). The most common cephalopod prey item was the Common Octopus (*Octopus vulgaris*), which was present in 32.91% of the samples, followed by the pencil squids (family Loliginidae), present in 31.65%. However, in terms of number of reads, pencil squids comprised 7.68% of all reads and octopodids only 4.46%. Finally, Oegopsida squids (Family Histiotheutidae) were present in 2.53% of the samples but comprised less than 1% of the reads (Table 2; Fig. 2).

### Seasonal variation in the diet of *<u>P.westlandica</u>*

According to the Frequency of Occurrence (FOO) and the Relative Read Abundance (RRA), our results show differences between seasons (Fig. 3) and between sampling sites (Fig.4).

**Figure 3.**
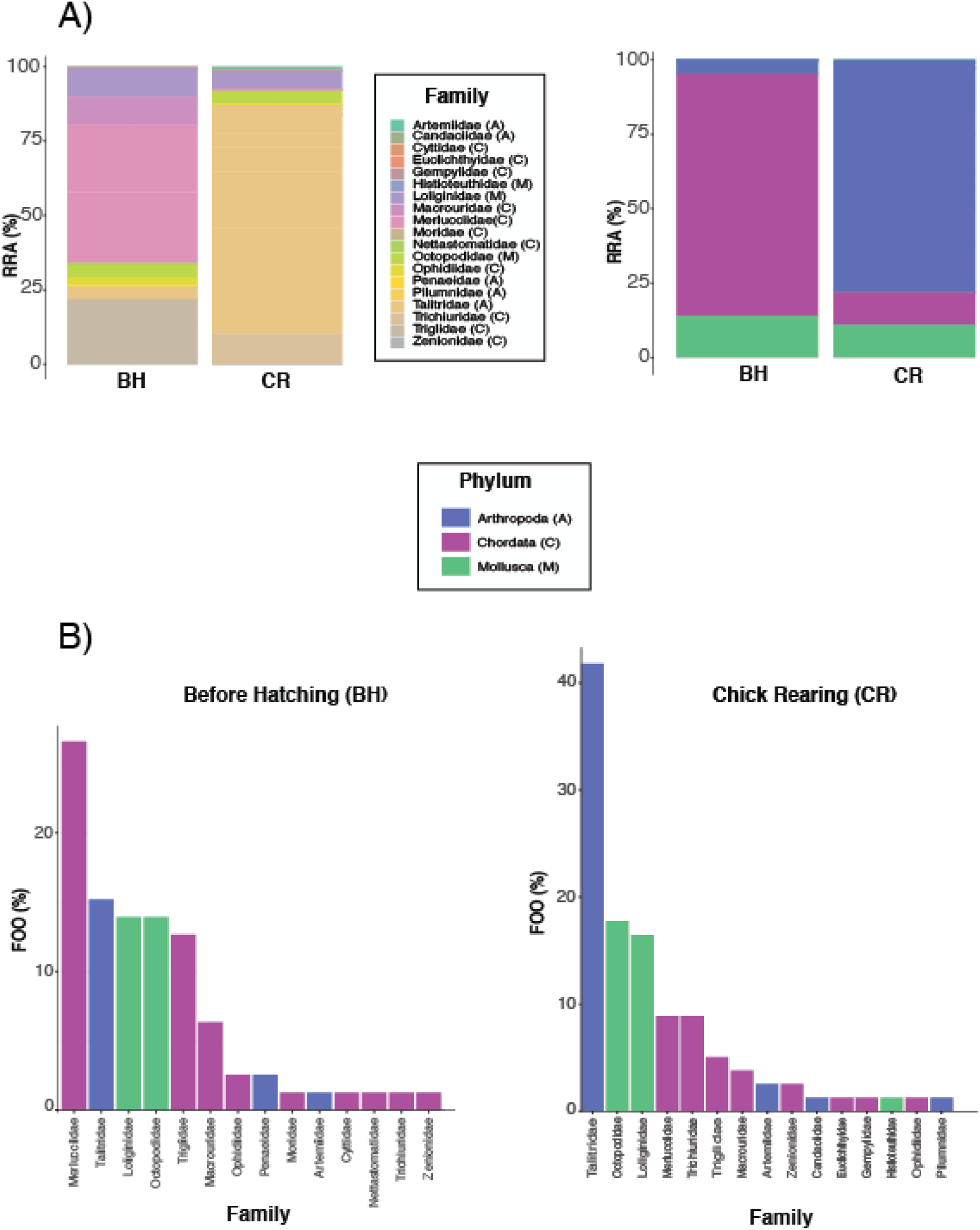
Seasonal variations at family level between the early breeding season or before hatching (BH) and the late breeding season or chick-rearing (CR), according to two biodiversity metrics: A) Relative Read Abundance (RRA) and B) Frequency of Occurrence (FOO). Taxa with less than 1% of FOO or RRA were not included in the plots.

**Figure 4.**
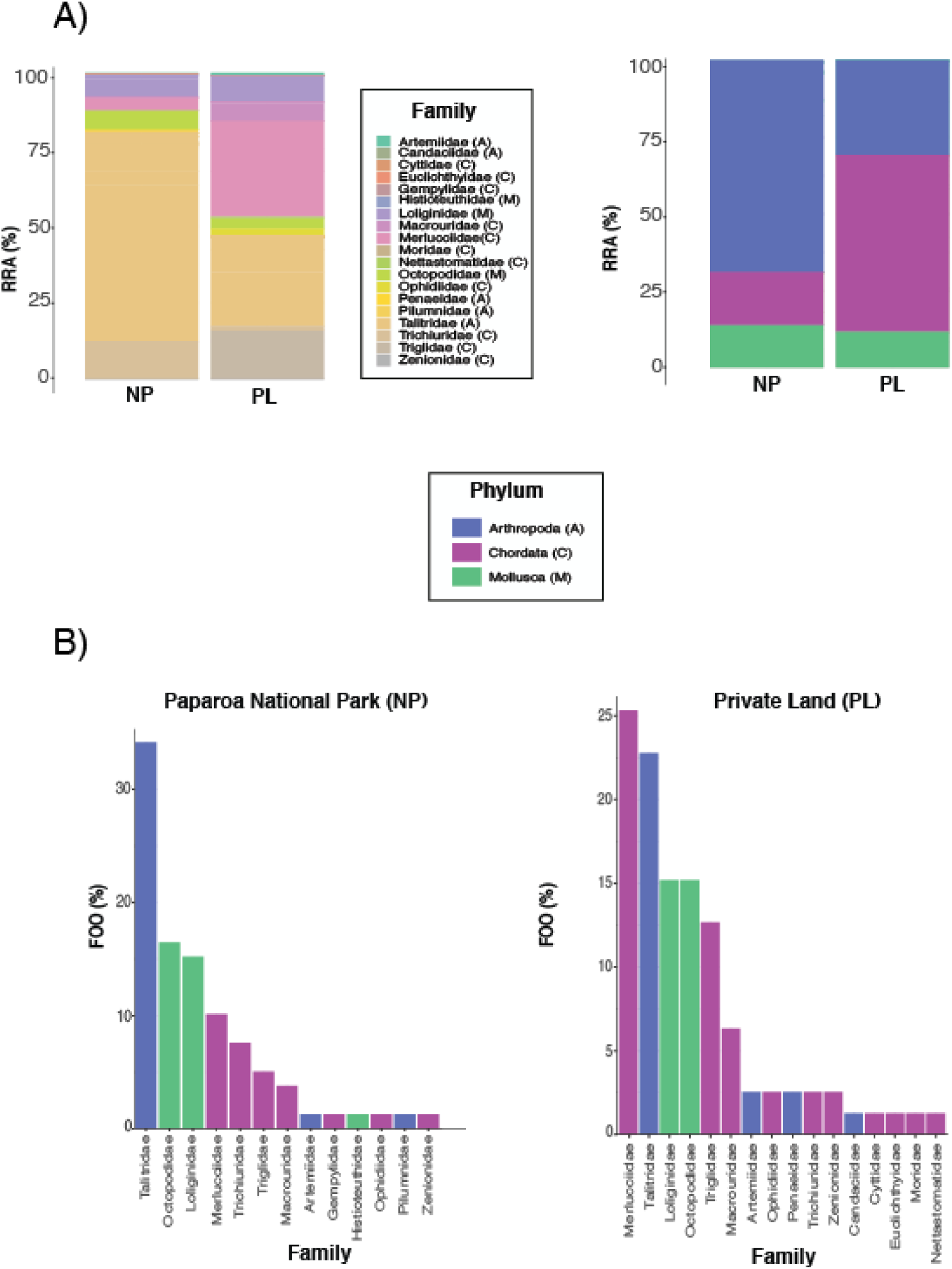
Geographical variations at family level between the two sub-colonies: the Paparoa National Park (NP) and the private land (PL), according to two biodiversity metrics: A) Relative Read Abundance (RRA) and B) Frequency of Occurrence (FOO). Taxa with less than 1% of FOO or RRA were not included in the plots.

Prey community composition varied significantly between the two different seasons both in terms of read abundance (Dev_1,79_ = 232.5, p = 0.004) and prey occurrence (Dev_1,79_ = 189.2, p = 0.004) (Fig. 3). These differences are clearly visible on the graphical ordination (Fig. S5).

When looking at frequency of occurrence, during the early breading season (before hatching), merluccids were the most common prey, followed by Talitroidea and then by cephalopods (pencil squids and octopuses showing the same value of FOO). In contrast, during the late breading season (chick rearing), Talitroidea were the most common prey followed by octopodids and pencil squids (Table 2; Fig.3A). A similar pattern was observed for relative read abundance, although with greater differences in the metric values (Table 2; Fig.3B).

Talitroidea were the most common prey group overall and during Chick-rearing (CR), representing more than 99% of all arthropods identified in this study. Although a minor prey, the Banana shrimp (*Penaeus merguiensis*), was present in 2.53% of samples before hatching but it was absent during the chick-rearing season. In the same way, the Bristly crab (*Pilumnus hirtellus*) and *Candacia armata* comprised both 1.27% of samples before hatching and were absent during chick-rearing (Table 2).

Fifteen OTUs of Actinopterygii fish were identified in the samples collected before hatching (13 identified at species level and 2 at family level), compared to 9 OTUs (corresponding to 8 species) during the chick-rearing season. Hoki was the most common fish species detected before hatching, followed by Cocky gurnard and Southern hake. During the chick-rearing season, Trichiuridae fish were the most common followed by Southern hake and Cocky gurnard (both showed the same FOO value).

With regards to cephalopods, Pencil squids (Loliginidae) and octopodids (Octopodidae) were present in the same number of samples, while, during the chick-rearing season, octopodids were more common than pencil squids. Interestingly, an Oegopsida squid (Histioteuthidae) was also detected during the chick-rearing season while it was completely absent before hatching (Table 2; Fig.3A and B).

Regarding species richness, the values of alpha diversity (Simpson) were not significantly different between seasons, with before hatching **α** = 0.31 ± 0.05 [mean ± SE] chick-rearing season **α** = 0.28 ± 0.04 [mean ± SE] (Fig.5).

**Figure 5.**
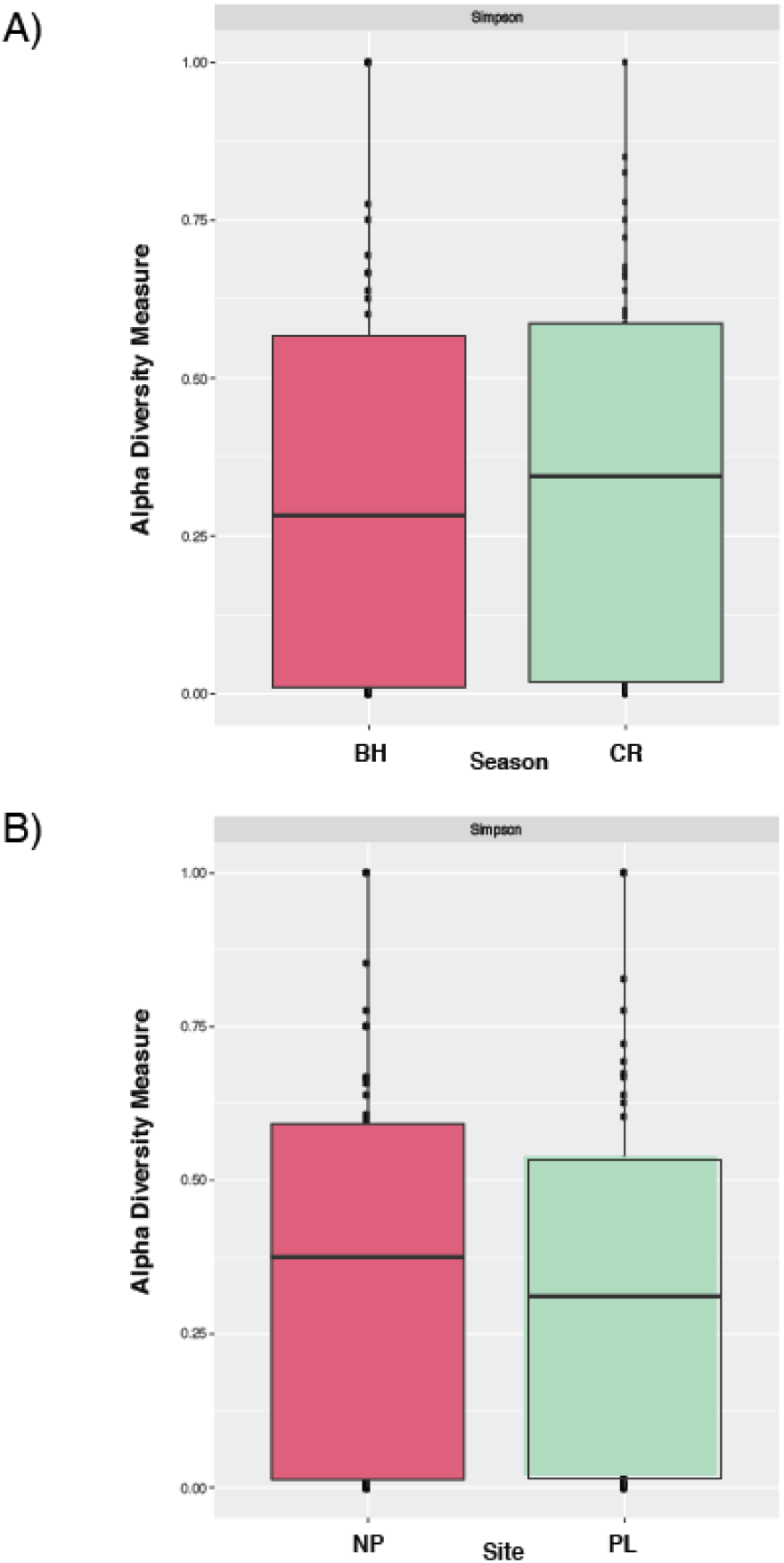
A) Seasonal and **B)** geographical differences in prey items according to alpha diversity measures.

### Geographical variation in the diet of *<u>P.westlandica</u>*

Significant differences in prey community were observed between the two sub-colonies, both in terms of read abundance (Dev_1,79_ = 203, p = 0.002) and occurrence of prey items (Dev_1,79_ = 172.6, p = 0.003) (Fig. 4).

Differences in prey community composition between sub-colonies are visible on the ordination biplot (Fig.S5).

Arthropods (Talitroidea) were found to be by far the most commonly detected prey group in the sub-colony located within the Paparoa National Park (NP), followed by octopodids and pencil squids. In contrast, in the Private Land (PL), merluccids were the most common group of prey, followed by Talitroidea and pencil squids.

Eleven OTUs of Actinopteriigy were identified in samples collected in PL (10 identified at species level and 1 at family level), while 17 OTUs (15 identified at species level and 2 at family level) were found in NP. Cutlassfish (family Trichiuridae) were the most common prey within NP, followed by Hoki, Southern hake and Cocky gurnard. In PL samples, however, Hoki was the most common fish taxa, followed, in this case, by Cocky gurnard and Southern hake (Table 2).

With regards to cephalopods, Common Octopuses were the most common group followed by pencil squids in NP, and both were present in the same number of samples in PL. In terms of read abundance, pencil squids were slightly more abundant than Common Octopuses in NP and in PL. (Table 2; Fig.4A and B).

Seasonal variation, no significant differences in species richness (alpha diversity) were observed in prey diversity when comparing the two sub-colonies NP (**α** [mean ± SE] = 0.31 ± 0.05) and PL (**α** [mean ± SE] = 0.28 ± 0.04) (Fig.5).

## Discussion

Our study is the first attempt to characterize the diet of the New Zealand endemic Westland petrel using DNA metabarcoding. By using DNA sourced from faecal material we were able to demonstrate how a non-invasive dietary DNA (dDNA) approach can be used to describe the diet of this endangered species. We found that amphipods were the most common prey, followed by cephalopods and fish. These results could correspond to natural foraging behaviour, but also support close links between Westland petrel diets and New Zealand’s commercial fishing activities. The high abundance of amphipods could be due to petrels feeding on discarded fisheries waste (fish guts). We also showed significant differences in diet between seasons (before hatching vs chick-rearing season) and between sampling sites (two sub-colonies 1.5 km apart), indicating that foraging strategies of the Westland petrel can be flexible. Our dDNA approach can contribute to the conservation of seabirds by non-invasively describing diet using metabarcoding.

Although metabarcoding has the potential to be an extremely valuable tool in conservation genomics, it still has deficiencies that need to be resolved. In our approach, we were able to infer 88.6% of the prey species within the diet of the Westland petrel, however, the resolution of the amplicon was insufficient for assigning Talitroidea to species level. This limitation may be due to the short size of the amplicon and/or the incompleteness of existing genetic databases (Gold et al., 2021; Hleap et al., 2021; Liu et al., 2020; Pompanon et al., 2012; Wangensteen et al., 2018). For this reason, we cannot confirm whether Talitroidea is a primary prey. In addition, our primer set could potentially have higher affinity with arthropods (or even amphipods) (Elbrecht and Leese, 2015) making Talitroidea overrepresented in the characterization of the diet of the Westland petrel. The two primer pairs approach did not provide any valuable information in our case as the Chordata primer pair did not add any valuable information to the data set obtained with the Malacostraca primer pair. However, overall we were able to infer the diet of this endangered seabird, have an insight into its ecological network and identify key prey species for the survival of Westland petrels.

Previous works on the diet of *P. westlandica* were based on morphological identification of prey remains, and carried out exclusively during the breeding or chick-rearing season (Freeman, 1998; Imber, 1976). The observed seasonal and geographical variations in the diet of *P. westlandica* provide a broad picture of the feeding requirements and foraging ecology of this species. Our study shows the presence of fish, cephalopods and amphipods (crustaceans) in the diet of *P. westlandica*, confirming the results of previous approaches (Freeman, 1998; Imber, 1976). However, the relative importance of each type of prey differs considerably between these studies and the current work, where we identified a number of taxa undetected before in such high proportions.

The phylum showing the highest percentage of prey reads was Arthropoda (45.57% of the reads, compared to 42.14% of the reads for fish). Arthropoda reads were mainly represented by Talitroidea (landhoppers or sandhoppers) (order Amphipoda). With this approach we cannot guarantee that these animals, ranging from 1 mm to 34 cm in size, are primary or secondary prey of the Westland petrel. Most species are microscopical benthic zooplankton and are known to be common prey of many cephalopods (Villanueva et al., 2017) and fish, including Hoki (Connell et al., 2010; Livingston and Rutherford, 1988) and Hake (Dunn et al., 2010). Therefore, Amphipods detected in this study could potentially be secondary prey. On the other hand, it is known that several Procellariiformes feed within coastal areas, which is the environment where amphipods are more present and reachable for seabirds (Thomas et al., 2006; Warham, 1996). Moreover, several seabirds such as penguins feed regularly on amphipods (Jarman et al., 2013; Knox, 2006), and large amphipods could potentially represent a fundamental food source for Antarctic seabirds (Centro de Investigacion Dinamica de Ecosistemas Marinos de Altas Latitudes, 2017), where they play a similar role as the krill (Euphausiacea) in the water column. Moreover, amphipods are found in the stomachs of other Procellariiformes, such as the Providence petrel (*Pterodroma solandri*) (Bester et al., 2011; Lock et al., 1992), the Blue petrel (*Halobaena caerulea*) (Croxall, 1987) and the Wilson’s storm petrel (*Oceanites oceanicus*). These birds are known to feed on amphipods when krill is not available (Quillfeldt et al., 2019, 2005, 2001, 2000). Imber (1976) found no planktonic crustacean in the stomach of *P. westlandica* and Freeman (1998) only detected small percentage of taxa belonging to three different families: Euphausiidae or krill (*Nyctiphanes australis* and *Thysanoessa gregaria*), Caridea or caridean shrimps (*Notostomus auriculatus* and an unidentified species) and Cymothoidae (unidentified species). Another possible explanation lies in the geographic distribution of arctic benthos, including amphipods, which is now displaying a hotspot in the south of New Zealand due to the climate change (Barnes et al., 2009). This potential increase in abundance could have increased the availability of amphipods for the petrel. In short, it still remains unclear whether Amphipods are primary prey, secondary prey (Sheppard et al., 2005) or both, but we can confirm that these taxa play a major role in the flow of energy through the food web. Further research, potentially using a food web approach in which diets from each of the components of the network are characterized, would be useful to fill this gap of knowledge.

Fish are major prey items of Procellariiformes (Bester et al., 2011; Bocher et al., 2000; da Silva Fonseca and Petry, 2007; Freeman, 1998; Imber, 1976; Prince and Morgan, 1987; Spear et al., 2007; Stewart et al., 1999), and the Westland petrel is not an exception. According to our results, fish (all belonging to the order Actinopteriigy) represent 15.03% of the prey reads, and they are the second most important phylum, in terms of RRA. In addition, fish DNA was detected in 37.93% of the samples. The fish species identified by our approach are consistent with previous studies (Freeman, 1998; Imber, 1976) but also include new species. In concordance with previous knowledge, the Hoki was identified as the most abundant fish prey item. However, we also found Hake, another Merlucciidae, and Cocky gurnard (which was not identified by previous approaches), followed by Hoki in abundance and occurrence,.

Hoki and Hake live between 28 and 1,000 m below sea level (Table 1), which makes these fish rarely catchable naturally for Westland petrels, since the birds can only dive down to 15 m below the surface (Freeman, 1998). However, these species, especially Hoki, are some of the main fishery species caught in New Zealand waters (Livingston and Rutherford, 1988). The fishing season for Merlucciids spans mainly between June and September, thereby encompassing most of the Westland petrel’s breeding season (Waugh et al., 2018; Waugh and Wilson, 2017), and including both sampling events of this study. Thus, the Westland petrels could scavenge these fish species from fishing vessels. In many cases, what is available for seabirds in the fishing boat decks are the leftovers from the fish, such as stomachs. These stomachs may contain fish prey items, which could explain the high abundance of Talitroidea in our results.

The same conclusion could apply to a number of other fish species with deep depth ranges, that are naturally unreachable for the petrel, but are important fishery species (Freeman, 1998; Froese and Pauly, 2010). These include rattails (Macrouridae), such as the Thorntooth grenadier as well as two newly identified prey items, namely the Hacknose grenadier and the Banded whiptail, among other fish species living in deep sea waters (Table 1). In the case of Hoki, however, natural predation may also be possible at night, as this fish species is known to migrate to surface waters to feed during the night (McClatchie and Dunford, 2003; O’Driscoll et al., 2009), when *P. westlandica* forages more actively (Waugh et al., 2018).

Cocky gurnard, which can sometimes be found in shallow waters (Froese and Pauly, 2010), could be caught naturally by the petrel. However, as stated before, it is also a known fishery species that could have been scavenged from the fishing waste. In addition, many fish species belonging to the family Trichiuridae can live close to the surface. Myctophid fishes, which were reported to be natural prey of the Westland petrel (Freeman, 1998; Imber, 1976), were not identified in our sampling. It is possible that these species are no longer available or no longer selected by the Westland petrel, as previous studies were conducted more than 20 years ago for Freeman (Freeman, 1998) and more than 45 years ago for (Imber, 1976).

In any of the potential fishing scenarios (natural fishing or scavenging) this study confirms the importance of fish prey items in the diet of the petrel, which could extensively use fish waste from the Hoki fishery and other inshore small fisheries, at least in the winter season (Freeman, 1998), but they could also catch some fish species naturally in certain situations. It is common for opportunistic seabirds to feed on fishery waste, however, if the dependence on this food source is very high, changes and new regulations in fishing activity could modify the birds’ behaviour and potentially impact their survival and population size (Abrams, 1983; Freeman, 1998; Oro et al., 1996, 1995). We identified Hoki, Southern Hake and Cocky gurnard as key prey species for the Westland petrel. Thus, wild populations of these fish species and fishing activities should be managed in a way that maintain these resources available for the petrel.

Cephalopods are also a key component of the diet of the Westland petrel as they comprised 12.29% of prey reads, and these taxa were detected in 53.16% of the samples. Six out of eight cephalopod OTUs could only be assigned to family level. Only the common Octopus (*Octopus vulgaris*) was assigned to species level, a taxon already found in previous studies (Freeman, 1998; Imber, 1976). Our results are consistent with Freeman (1998), which states that fish prey items are followed by cephalopods within the Westland petrels’ diet. In the case of *Histioteuthis sp.,* they are deep-sea squid (Voss et al., 1998), but migrate to surface waters at night by vertical migration (Roper and Young, 1975), which makes them catchable by Westland petrel. The other two families, Loliginidae and Octopodidae (Common octopus), which were also identified in previous studies, are present from surface waters down to 500 m deep, and thus naturally catchable for the Westland petrel. Nevertheless, these families also include several commercial species as well as species commonly reported as bycatch (Davies et al., 2009; Pierce et al., 2010). Therefore, it is possible that petrels fed on some cephalopods through fishery waste.

A number of other Mollusca prey species were, listed in previous studies (Freeman, 1998; Imber, 1976), but not detected in our approach. These include cephalopods belonging to the orders Sepioidea or Vampyromorpha, among others. It is unclear whether their absence in our analysis is due to the lack of genetic sequences in the NCBI database or a change in the feeding habits of the birds in the past 20 years. Further research focusing on Mollusca would be required to solve this gap of knowledge.

Marked dietary switches between breeding and non-breeding seasons have been documented for several seabirds (Howells et al., 2018), and are considered a sign of plasticity in behaviour (Quillfeldt et al., 2019). These switches may reflect variation in prey availability, a change of strategy between seasons, or a combination of both (Howells et al., 2018; Paleczny et al., 2015; Sydeman et al., 2015). Because these variations can severely affect populations of marine top predators (Cury et al., 2000; Reid and Croxall, 2001) it is essential to understand their drivers to ensure the conservation of the Westland petrel. Adaptability to different temperatures and availability of resources would be a sign of resilience of the petrels’ populations to different environments and can greatly inform the design of conservation plans (Berkes and Jolly, 2002; Jones et al., 2020; McDonnell and Hahs, 2015; Yellen, 1977).

As hypothesized, we found a clear seasonal variation in the diet of *P. westlandica*, both in terms of read abundance and the occurrence of prey species, meaning that the composition of the diet changes in a substantial way between incubation and chick-rearing season. This change is particularly visible for fish (specifically merluccids) and Talitroidea, with fish being the most abundant prey before hatching while Talitroidea are by far the most common prey during the chick-rearing season. One explanation could be that adult petrels feed their chicks with highly nutritive fish and cephalopods, while they feed themselves mainly with crustaceans (and some cephalopods). This hypothesis is highly consistent with the significant loss of weight in adult seabirds during the breeding season, while their chicks experience rapid growth (Ainley, 1990; Barrett et al., 1985; Leal et al., 2017). In this case, the choice of prey items by adults may be influenced by the developmental stage and the needs of the chicks. Despite these seasonal differences in prey preferences, prey species richness remains similar between in seasons.

Our results suggest that seasonal variations may be more influenced by changes in foraging strategy, rather than changes in prey availability. Indeed, the peak of the Hoki fishery in New Zealand encompasses both July (before hatching period) and September (chick-rearing period), which means, fishery waste would be equally available during both seasons. Such changes in foraging strategy reflect an adaptation to new conditions and environments, which ensures that petrels can find suitable feeding resource throughout the breeding season.

Regarding sub-colonies and contrary to our expectation, we found significant differences in prey composition between both sub-colonies. A possible explanation of these differences could be that seabirds from nearby sub-colonies forage in different locations, possibly to avoid or decrease inter-colony competition (Cecere et al., 2015; Grémillet et al., 2004; Wakefield et al., 2013). Birds’ diet could also change every day depending on resource availability, and prey resources may have been very different in the two consecutive days used for collecting samples in both sub-colonies due to short-term variations in temperature and/or resource availability. Finally, the sub-colonies might be different genetic haplotypes, occupying slightly different dietary niches. In order to clarify the origin of these differences in prey community composition between sub-colonies, further studies on the foraging ecology and population genetics of the Westland petrel should be conducted.

Sustainable management of worldwide fishery industry needs information regarding the overlap of marine organisms, such as seabirds, with fishing industry (Frederiksen et al., 2004; McInnes et al., 2017b; Okes et al., 2009). Seabirds scavenge food from fishery waste, which results in a high number of incidental kills through bycatch, potentially disturbing population dynamics (Brothers, 1999; McInnes et al., 2017b; Sullivan et al., 2006; Tuck et al., 2011; Watkins et al., 2008; Waugh et al., 2008; Waugh and Wilson, 2017). Yet, the diet of seabirds relies on this commercial activity, as fishery waste represents a nutritious food source, naturally unreachable for seabirds. Therefore, understanding these interactions is essential for seabird conservation and efficient ecosystem-based fishing regulation (Becker and Beissinger, 2006; Freeman, 1998; Furness, 2003; Furness and Tasker, 2000; McInnes et al., 2017b; Phillips et al., 1999; Waugh et al., 2008). In this context, non-invasive dietary studies can provide knowledge to assess risks as well as the needs of these species that may rely heavily on commercial fishing activity (Gaglio et al., 2018; McInnes et al., 2017a, 2017b). This issue is particularly urgent in the case of endangered species, such as the Westland petrel, and, in this study, we show a probable link between fisheries in New Zealand and the diet of the petrel, that should be considered in management and conservation strategies.

Our results show the potential of non-invasive dietary studies in highlighting the reliance of endangered seabirds on commercial fishing activity (Gaglio et al., 2018; McInnes et al., 2017a, 2017b). Such study should draw attention to the complexity that lies in the implementation of fishing regulations and the associated risks for the conservation of endangered species. In the case of Westland petrel, these regulations should take into account the close link between the commercial fishing and the diet preferences of the birds regarding fish and cephalopods. Several mitigation solutions have been suggested by practitioners or already included in conservation reports (OpenSeas, 2019), to limit the number of accidental kills in seabirds and find a sustainable equilibrium between fishery industry and threatened species. Hence, knowledge on how seabirds in general, and Westland petrel in particular, interact with fishing vessels and fishing gear is necessary to develop bycatch reduction techniques and using or developing gear less dangerous for the seabirds.

### Data accessibility

Data, bioinformatic scripts and R code for statistical analyses are available at https://figshare.com/s/9c6d1292b51d35daf422

## Supplementary material

**Table S1.** Sample list showing the sample identification code (ID), the season when it was collected and the exact date as well as the site where it was collected from.

**Figure S1.** Bioinformatic results from the 16S dietary metabarcoding approach showing the number of reads at each step of the filtering process.

**Figure S2.** Accumulation curve representing the cumulative number of prey OTUs detected against the number of faecal samples analysed (n = 87). Horizontal solid line represents the number of prey OTUs expected with limitless sampling, based on bootstrapped estimates.

**Figure S3.** Point plot representing the descending number of sequence reads per OTU detected.

**Figure S4.** Cumulative line plot of the frequency of OTUs detected per number of sequence reads (sequence depth).

**Figure S5.** Ordination biplot to visualize the differences in community composition of the diet of the Westland petrel. The different colours show the differences between the seasons (early -BH- and late breading season -CR-) and the shapes represent the different sites or sub-colonies (Natural Park -NP- and Private Land -PL-).

## Author contribution

Designed the study: SB. Obtained funding: SB. Collected samples: SB. Performed laboratory analyses: MCL, SB. Analysed the data and prepared the figures: MQ. Wrote the first draft of the manuscript MQ, SB. All authors contributed to the writing of the final manuscript.

## Acknowledgements

This study was funded by an internal Research Development Fund obtained by SB in 2015 at Unitec Institute of Technology (RI15012). We thank Susan Waugh from Office of the Parliamentary Commissioner for the Environment for providing early samples that were used for proof of concept, and for her comments and advice on a previous version of the manuscript. We are grateful to Conservation Volunteers NZ https://conservationvolunteers.co.nz/) and particularly James Washer for providing information about colony location outside of the protected area and logistical support on site. We thank Bruce Menteath from Petrel Colony tours (http://www.petrelcolonytours.co.nz/) for giving us access to the colony on his land and sharing his knowledge about the birds.

We also thank Louise Burkett and Amy Hou for their technical help as part of their internships at Unitec Institute of Technology. Also, we would like to thank Joan Garcia-Porta from Washington University of St. Louis (Missouri) for his help and advice in bioinformatics, David Ochoa Castañon for his helpful comments on bird behaviour and Lucas Sire for his help in community composition analyses.

Version 4 of this preprint has been peer-reviewed and recommended by Peer Community In Ecology (https://doi.org/10.24072/pci.ecology.100090).

