## Supplementary Material for "Metabarcoding faecal samples to investigate spatiotemporal variation in the diet of the endangered Westland petrel (*Procellaria westlandica*)"

**Investigating spatiotemporal variation in the diet of Westland Petrel through metabarcoding, a non-invasive technique**

Marina Querejeta^1^, Marie-Caroline Lefort^2,3^, Vincent Bretagnolle^4^, Stéphane Boyer^1,2^

^1^ Institut de Recherche sur la Biologie de l'Insecte, UMR 7261, CNRS-Université de Tours, Tours, France

^2^ Environmental and Animal Sciences, Unitec Institute of Technology, 139 Carrington Road, Mt Albert, Auckland 1025, New Zealand

^3^ Cellule de Valorisation Pédagogique, Université de Tours, 60 rue du Plat d’Étain, 37000 Tours, France.

^4^ Centre d’Études Biologiques de Chizé, UMR 7372, CNRS & La Rochelle Université, 79360 Villiers en Bois, France.

**Table S1**. Sample list showing the sample identification code (ID), the season when it was collected and the exact date as well as the site where it was collected from.

| **Sample ID** | **Season** | **Collection date** | **Site** |
| --- | --- | --- | --- |
| CV1 | Before Hatching | 09/07/2015-10/07/2015 | Paparoa Natural Park |
| CV2 | Before Hatching | 09/07/2015-10/07/2015 | Paparoa Natural Park |
| CV3 | Before Hatching | 09/07/2015-10/07/2015 | Paparoa Natural Park |
| CV4 | Before Hatching | 09/07/2015-10/07/2015 | Paparoa Natural Park |
| CV5 | Before Hatching | 09/07/2015-10/07/2015 | Paparoa Natural Park |
| CV6 | Before Hatching | 09/07/2015-10/07/2015 | Paparoa Natural Park |
| CV7 | Before Hatching | 09/07/2015-10/07/2015 | Paparoa Natural Park |
| CV8 | Before Hatching | 09/07/2015-10/07/2015 | Paparoa Natural Park |
| CV9 | Before Hatching | 09/07/2015-10/07/2015 | Paparoa Natural Park |
| CV10 | Before Hatching | 09/07/2015-10/07/2015 | Paparoa Natural Park |
| CV11 | Before Hatching | 09/07/2015-10/07/2015 | Paparoa Natural Park |
| CV12 | Before Hatching | 09/07/2015-10/07/2015 | Paparoa Natural Park |
| CV13 | Before Hatching | 09/07/2015-10/07/2015 | Paparoa Natural Park |
| CV14 | Before Hatching | 09/07/2015-10/07/2015 | Paparoa Natural Park |
| CV15 | Before Hatching | 09/07/2015-10/07/2015 | Paparoa Natural Park |
| CV16 | Before Hatching | 09/07/2015-10/07/2015 | Paparoa Natural Park |
| CV17 | Before Hatching | 09/07/2015-10/07/2015 | Paparoa Natural Park |
| CV18 | Before Hatching | 09/07/2015-10/07/2015 | Paparoa Natural Park |
| CV19 | Before Hatching | 09/07/2015-10/07/2015 | Paparoa Natural Park |
| CV20 | Before Hatching | 09/07/2015-10/07/2015 | Paparoa Natural Park |
| CV21 | Before Hatching | 09/07/2015-10/07/2015 | Paparoa Natural Park |
| CV22 | Before Hatching | 09/07/2015-10/07/2015 | Paparoa Natural Park |
| M1 | Before Hatching | 09/07/2015-10/07/2015 | Private Land |
| M2 | Before Hatching | 09/07/2015-10/07/2015 | Private Land |
| M3 | Before Hatching | 09/07/2015-10/07/2015 | Private Land |
| M4 | Before Hatching | 09/07/2015-10/07/2015 | Private Land |
| M5 | Before Hatching | 09/07/2015-10/07/2015 | Private Land |
| M6 | Before Hatching | 09/07/2015-10/07/2015 | Private Land |
| M7 | Before Hatching | 09/07/2015-10/07/2015 | Private Land |
| M8 | Before Hatching | 09/07/2015-10/07/2015 | Private Land |
| M9 | Before Hatching | 09/07/2015-10/07/2015 | Private Land |
| M10 | Before Hatching | 09/07/2015-10/07/2015 | Private Land |
| M11 | Before Hatching | 09/07/2015-10/07/2015 | Private Land |
| M12 | Before Hatching | 09/07/2015-10/07/2015 | Private Land |
| M13 | Before Hatching | 09/07/2015-10/07/2015 | Private Land |
| M14 | Before Hatching | 09/07/2015-10/07/2015 | Private Land |
| M15 | Before Hatching | 09/07/2015-10/07/2015 | Private Land |
| M16 | Before Hatching | 09/07/2015-10/07/2015 | Private Land |
| M17 | Before Hatching | 09/07/2015-10/07/2015 | Private Land |
| M18 | Before Hatching | 09/07/2015-10/07/2015 | Private Land |
| M19 | Before Hatching | 09/07/2015-10/07/2015 | Private Land |
| M20 | Before Hatching | 09/07/2015-10/07/2015 | Private Land |
| M21 | Before Hatching | 09/07/2015-10/07/2015 | Private Land |
| M22 | Before Hatching | 09/07/2015-10/07/2015 | Private Land |
| M23 | Before Hatching | 09/07/2015-10/07/2015 | Private Land |
| M24 | Before Hatching | 09/07/2015-10/07/2015 | Private Land |
| M25 | Before Hatching | 09/07/2015-10/07/2015 | Private Land |
| M26 | Before Hatching | 09/07/2015-10/07/2015 | Private Land |
| SCV1 | Chick Rearing | 22/09/2015-23/09/2015 | Paparoa Natural Park |
| SCV2 | Chick Rearing | 22/09/2015-23/09/2015 | Paparoa Natural Park |
| SCV3 | Chick Rearing | 22/09/2015-23/09/2015 | Paparoa Natural Park |
| SCV4 | Chick Rearing | 22/09/2015-23/09/2015 | Paparoa Natural Park |
| SCV5 | Chick Rearing | 22/09/2015-23/09/2015 | Paparoa Natural Park |
| SCV6 | Chick Rearing | 22/09/2015-23/09/2015 | Paparoa Natural Park |
| SCV7 | Chick Rearing | 22/09/2015-23/09/2015 | Paparoa Natural Park |
| SCV8 | Chick Rearing | 22/09/2015-23/09/2015 | Paparoa Natural Park |
| SCV9 | Chick Rearing | 22/09/2015-23/09/2015 | Paparoa Natural Park |
| SCV10 | Chick Rearing | 22/09/2015-23/09/2015 | Paparoa Natural Park |
| SCV11 | Chick Rearing | 22/09/2015-23/09/2015 | Paparoa Natural Park |
| SCV12 | Chick Rearing | 22/09/2015-23/09/2015 | Paparoa Natural Park |
| SCV13 | Chick Rearing | 22/09/2015-23/09/2015 | Paparoa Natural Park |
| SCV14 | Chick Rearing | 22/09/2015-23/09/2015 | Paparoa Natural Park |
| SCV15 | Chick Rearing | 22/09/2015-23/09/2015 | Paparoa Natural Park |
| SCV16 | Chick Rearing | 22/09/2015-23/09/2015 | Paparoa Natural Park |
| SCV17 | Chick Rearing | 22/09/2015-23/09/2015 | Paparoa Natural Park |
| SCV18 | Chick Rearing | 22/09/2015-23/09/2015 | Paparoa Natural Park |
| SCV19 | Chick Rearing | 22/09/2015-23/09/2015 | Paparoa Natural Park |
| SCV20 | Chick Rearing | 22/09/2015-23/09/2015 | Paparoa Natural Park |
| SCV21 | Chick Rearing | 22/09/2015-23/09/2015 | Paparoa Natural Park |
| SCV22 | Chick Rearing | 22/09/2015-23/09/2015 | Paparoa Natural Park |
| SCV23 | Chick Rearing | 22/09/2015-23/09/2015 | Paparoa Natural Park |
| SCV24 | Chick Rearing | 22/09/2015-23/09/2015 | Paparoa Natural Park |
| SCV25 | Chick Rearing | 22/09/2015-23/09/2015 | Paparoa Natural Park |
| SCV26 | Chick Rearing | 22/09/2015-23/09/2015 | Paparoa Natural Park |
| SCV27 | Chick Rearing | 22/09/2015-23/09/2015 | Paparoa Natural Park |
| SM1 | Chick Rearing | 22/09/2015-23/09/2015 | Private Land |
| SM2 | Chick Rearing | 22/09/2015-23/09/2015 | Private Land |
| SM3 | Chick Rearing | 22/09/2015-23/09/2015 | Private Land |
| SM4 | Chick Rearing | 22/09/2015-23/09/2015 | Private Land |
| SM5 | Chick Rearing | 22/09/2015-23/09/2015 | Private Land |
| SM6 | Chick Rearing | 22/09/2015-23/09/2015 | Private Land |
| SM7 | Chick Rearing | 22/09/2015-23/09/2015 | Private Land |
| SM8 | Chick Rearing | 22/09/2015-23/09/2015 | Private Land |
| SM9 | Chick Rearing | 22/09/2015-23/09/2015 | Private Land |
| SM10 | Chick Rearing | 22/09/2015-23/09/2015 | Private Land |
| SM11 | Chick Rearing | 22/09/2015-23/09/2015 | Private Land |
| SM12 | Chick Rearing | 22/09/2015-23/09/2015 | Private Land |
| SM13 | Chick Rearing | 22/09/2015-23/09/2015 | Private Land |
| SM14 | Chick Rearing | 22/09/2015-23/09/2015 | Private Land |
| SM15 | Chick Rearing | 22/09/2015-23/09/2015 | Private Land |
| SM16 | Chick Rearing | 22/09/2015-23/09/2015 | Private Land |
| SM17 | Chick Rearing | 22/09/2015-23/09/2015 | Private Land |
| SM18 | Chick Rearing | 22/09/2015-23/09/2015 | Private Land |
| SM19 | Chick Rearing | 22/09/2015-23/09/2015 | Private Land |
| SM20 | Chick Rearing | 22/09/2015-23/09/2015 | Private Land |
| SM21 | Chick Rearing | 22/09/2015-23/09/2015 | Private Land |
| SM22 | Chick Rearing | 22/09/2015-23/09/2015 | Private Land |
| SM23 | Chick Rearing | 22/09/2015-23/09/2015 | Private Land |
| SM24 | Chick Rearing | 22/09/2015-23/09/2015 | Private Land |

**Figure S1**. Bioinformatic results from the 16S dietary metabarcoding approach showing the number of reads at each step of the filtering process.

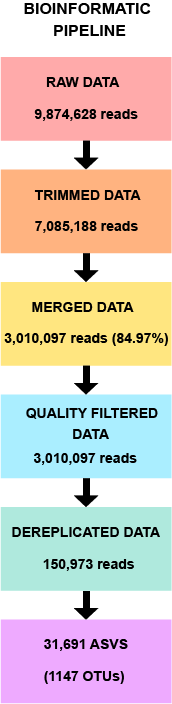

**Figure S2.** Accumulation curve representing the cumulative number of prey OTUs detected against the number of faecal samples analysed (n = 87). Horizontal solid line represents the number of prey OTUs expected with limitless sampling, based on bootstrapped estimates.

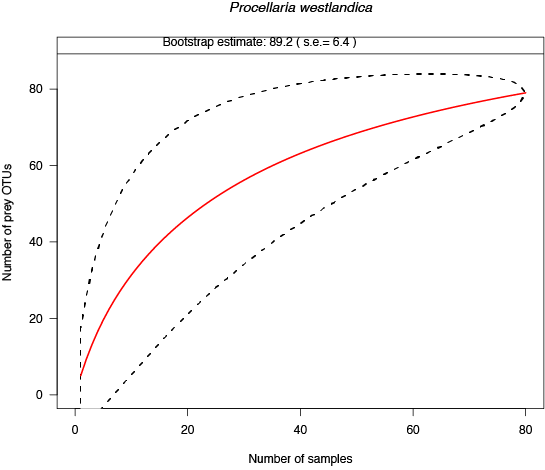

**Figure S3.** Point plot representing the descending number of sequence reads per OTU detected.

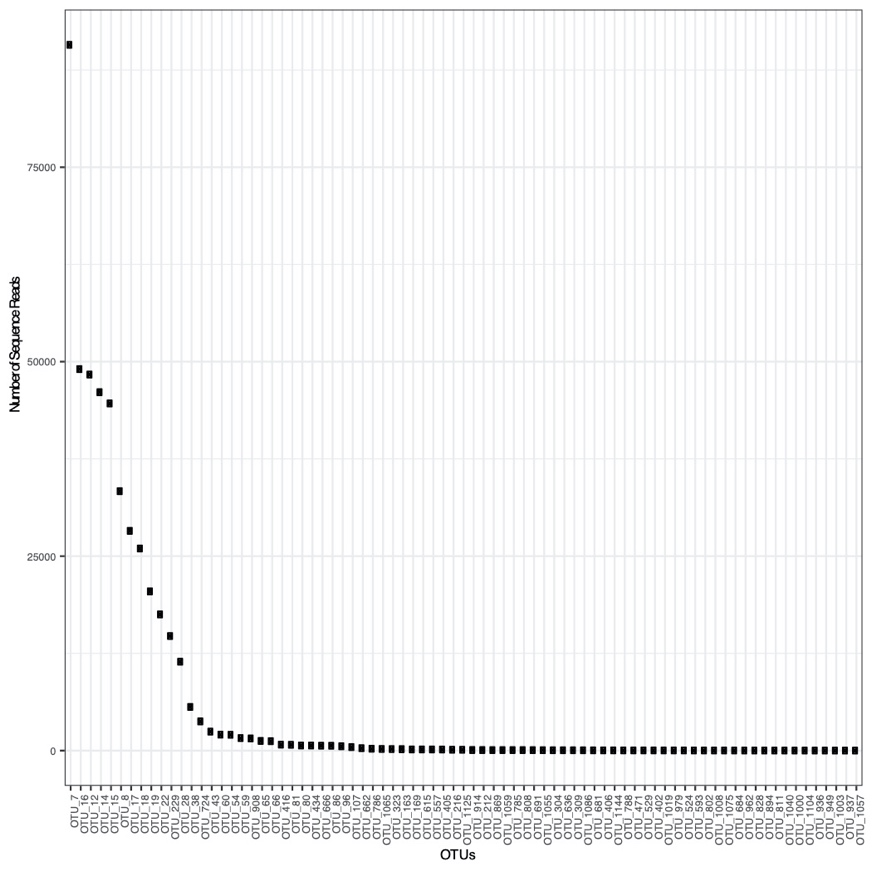

**Figure S4.** Cumulative line plot of the frequency of OTUs detected per number of sequence reads (sequence depth).

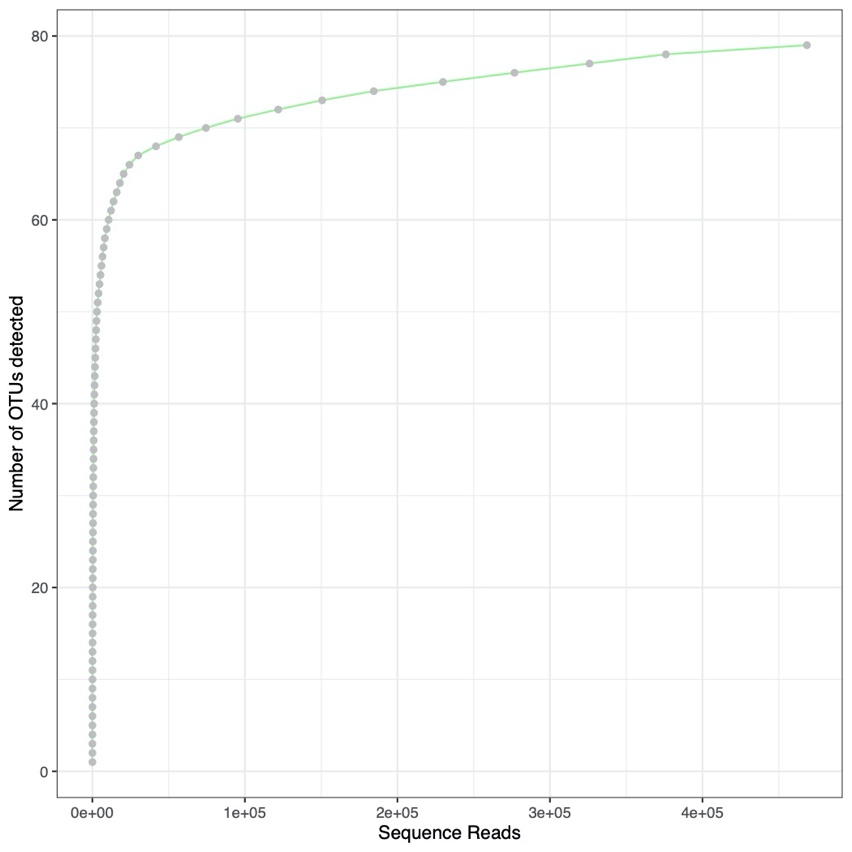

**Figure S5.** Ordination biplot to visualize the differences in community composition of the diet of the Westland petrel. The different colors show the differences between the seasons (early -BH- and late breading season -CR-) and the shapes represent the different sites or sub-colonies (Natural Park -NP- and Private Land -PL-).

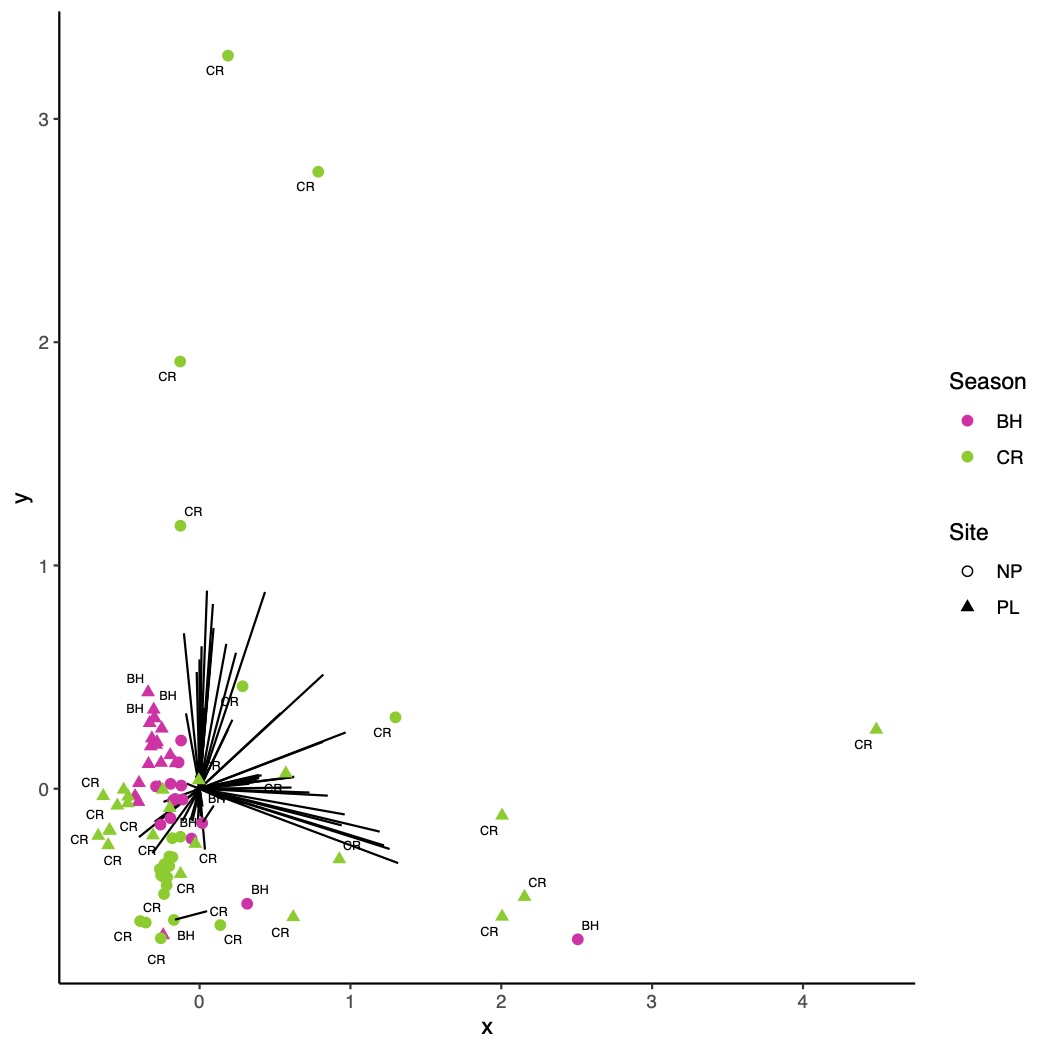
